# Parent-of-origin classification for diploid genomes using Pooch

**DOI:** 10.64898/2026.11.23.767630

**Authors:** Juhyun Kim, Brandon D. Pickett, Jillian Belluck, Nancy F. Hansen, Melissa A. Wilson, Human Pangenome Reference Consortium, Sergey Koren, Arang Rhie, Adam M. Phillippy

## Abstract

Advances in long-read sequencing have enabled the generation of near-complete human genomes at scale, but trio sequencing has typically been required to determine the parent of origin for each haplotype. Parent-of-origin information is important for studying inheritance patterns and imprinting-associated genetic disorders. Here we demonstrate that parent of origin can be reliably determined for all chromosomes of a Hi-C-phased diploid assembly using differentially methylated CpGs alone. Using 123 near-complete, trio-phased assemblies from the Human Pangenome Reference Consortium, we systematically cataloged differential methylation genome-wide, identifying 880 imprinted differentially methylated regions (iDMRs), including 150 with strong and 730 with moderate differential methylation, including two iDMRs on chrX. From these data we trained an elastic net logistic regression model (Pooch) that predicts parent of origin for each haplotype pair with a mean test accuracy of 0.997 and generalizes across sequencing technologies, cell types, and abnormal karyotypes. Notably, 91.7% of informative CpG features identified by the model were located outside of known iDMRs, suggesting that parent-of-origin differential methylation extends beyond previously established imprinted regions. We further identified parent-of-origin signals on chrX by correcting for the effects of X chromosome inactivation, revealing imprinted differentially methylated regions within an intron of *DHRSX* and in the promoter of *SRPK3*, where methylation depended on both parent of origin and X chromosome inactivation status. Our results show that it is now possible to achieve haplotype-resolved diploid genome assemblies routinely and determine maternal and paternal haplotypes for all chromosomes without the need for trios.

## Main

Imprinted differentially methylated regions (iDMRs) retain the methylation patterns inherited from egg and sperm and are a recurring biological phenomena^1,2^. Although most methylation marks are erased in the zygote after fertilization and re-established afterward, imprinted regions maintain their methylation levels throughout development^3^. These regions are enriched at the promoters of genes involved in growth, cognitive function, and essential fetal development^4–7^.

Disruption of these methylation patterns can lead to several diseases, such as Transient Neonatal Diabetes Mellitus Type 1^8^, Silver-Russell Syndrome^9^, and Beckwith-Wiedemann Syndrome^10,11^. Many parent-of-origin-specific methylation patterns are maintained throughout life in somatic tissues and are important for clinical genome interpretation.

iDMR discovery has typically relied on array-based platforms^12,13^ or short-read sequencing data^14^ mapped to a pseudo-haploid reference genome, mainly hg19 or GRCh38. However, these approaches have several limitations, including mapping errors and the incompleteness of the reference genome, which contains gaps and assembly errors^15,16^. This can overlook biologically important methylation patterns, especially with the repetitive regions of the genome^17^. To overcome these limitations, long-read sequencing^18–21^, complete genomes^17,22–24^ and pangenomes are increasingly being adopted^22,25^.

Classifying parent-of-origin directly from clinical samples, without the need for parental sequencing, would facilitate the investigation of imprinting effects and the potential causal roles of iDMRs in disease. However, assigning parental origin to assembled haplotypes remains challenging and typically requires parental data to identify parent-specific k-mers that distinguish maternal (Mat) and paternal (Pat) haplotypes^26,27^. It has been shown that parent-of-origin can be assigned for chromosome-scale haplotypes using previously established iDMRs^19^, but this level of phasing has previously required specialized techniques such as Strand-seq. Complete, “telomere-to-telomere” (T2T) genome assemblies have recently become more accessible thanks to the combination of long-read and Hi-C sequencing^28^. Importantly, long-read sequencing can detect methylated bases, while Hi-C enables chromosome-scale phasing. Such T2T genome assemblies are now being generated at increasing scale, including by the Human Pangenome Reference Consortium (HPRC)^29,30^ and T2T Consortium^23,31^, as well as from cancer genomes ^32–35^ and samples with unusual karyotypes^36^. Recent advances have also enabled high-quality assemblies using Nanopore sequencing alone^37–39^.

While the combination of these techniques has enabled parent-of-origin assignment without the need for trios, the X chromosome has remained elusive due to its lack of established iDMRs. In female eutherian mammals, one of the two X chromosomes is randomly selected for transcriptional silencing through the action of the long noncoding RNA X-inactive specific transcript (*XIST*), which initiates epigenetic modifications, including DNA methylation, to establish and maintain X chromosome inactivation (XCI)^40^. In contrast, marsupials exhibit imprinted XCI, in which the paternally inherited X chromosome is consistently inactivated throughout life^41^. Similar paternal XCI is observed during early embryonic development in mice; however, this imprint is subsequently erased and replaced by random XCI in the embryo proper^41–43^. Due to these complications, and the added difficulty of assembling human sex chromosomes, stable iDMRs have not yet been identified on human chrX.

Here, we present a comprehensive catalog of iDMRs derived from the HPRCv2 pangenome, defined here as any genomic region exhibiting differential methylation associated with the parent of origin. We also introduce Pooch (<u>P</u>arent-<u>o</u>f-<u>o</u>rigin <u>c</u>lassifier for <u>h</u>aplotypes), a method to predict parent of origin directly from differentially methylated CpGs. To facilitate the application of Pooch to new assemblies, species, and datasets, we provide an end-to-end pipeline for custom model training and inference. Pooch accurately determines parent-of-origin for all chromosome pairs, including chromosome X, without the need for trio data, and generalizes across cell types, sequencing technologies, and abnormal karyotypes. By accounting for XCI and its effect on methylation, we also revealed two regions near opposite ends of chromosome X whose methylation patterns depend on both parent of origin and XCI status.

## Results

### Genome-wide ab initio identification of imprinted differentially methylated regions

To identify a comprehensive set of iDMRs, we began with a set of 123 diploid assemblies from the HPRC for which each pair of chromosomes had been previously scaffolded and phased into maternal (Mat) and paternal (Pat) T2T haplotypes using short-read trio data (Supplementary Table 1). We aligned Oxford Nanopore (ONT) long reads from the same lymphoblastoid cell lines (LCLs) to their matched diploid assembly, called DNA methylated CpG sites, and lifted the resulting calls from each individual’s personalized genome to T2T-CHM13v2.0 coordinates. We then grouped methylation calls by haplotype and identified iDMRs based on differential methylation between the two haplotypes using DSS^44^ with a paired design to account for sample-specific variability and potential batch effects. Because the clonal LCLs do not show uniformly random X inactivation and to ensure a balanced representation of XCI patterns, we balanced the numbers of samples showing skewed maternal and paternal X-inactivation based on methylation of the *XIST* promoter (Supplementary Fig. 1 and Supplementary Table 2).

Because *XIST* is expressed exclusively from the inactive X chromosome and repressed on the active X chromosome, hypomethylation of its promoter indicates an inactive X (Xi), while hypermethylation indicates an active X (Xa).

Genome-wide, a total of 880 iDMRs were identified and classified into four categories based on the hypermethylated parental haplotype (maternal, miDMRs or paternal, piDMRs) and the magnitude of differential methylation, with strong iDMRs defined by an absolute methylation difference ≥0.15 and moderate iDMRs by a difference <0.15 (Fig. 1A, Supplementary Fig. 2, Supplementary Table 3). Notably, despite the absence of previously reported iDMRs on chrX, we identified one strong miDMR within an intron of *DHRSX* and one moderate piDMR near the centromere (Fig. 1A).

**Fig. 1:**
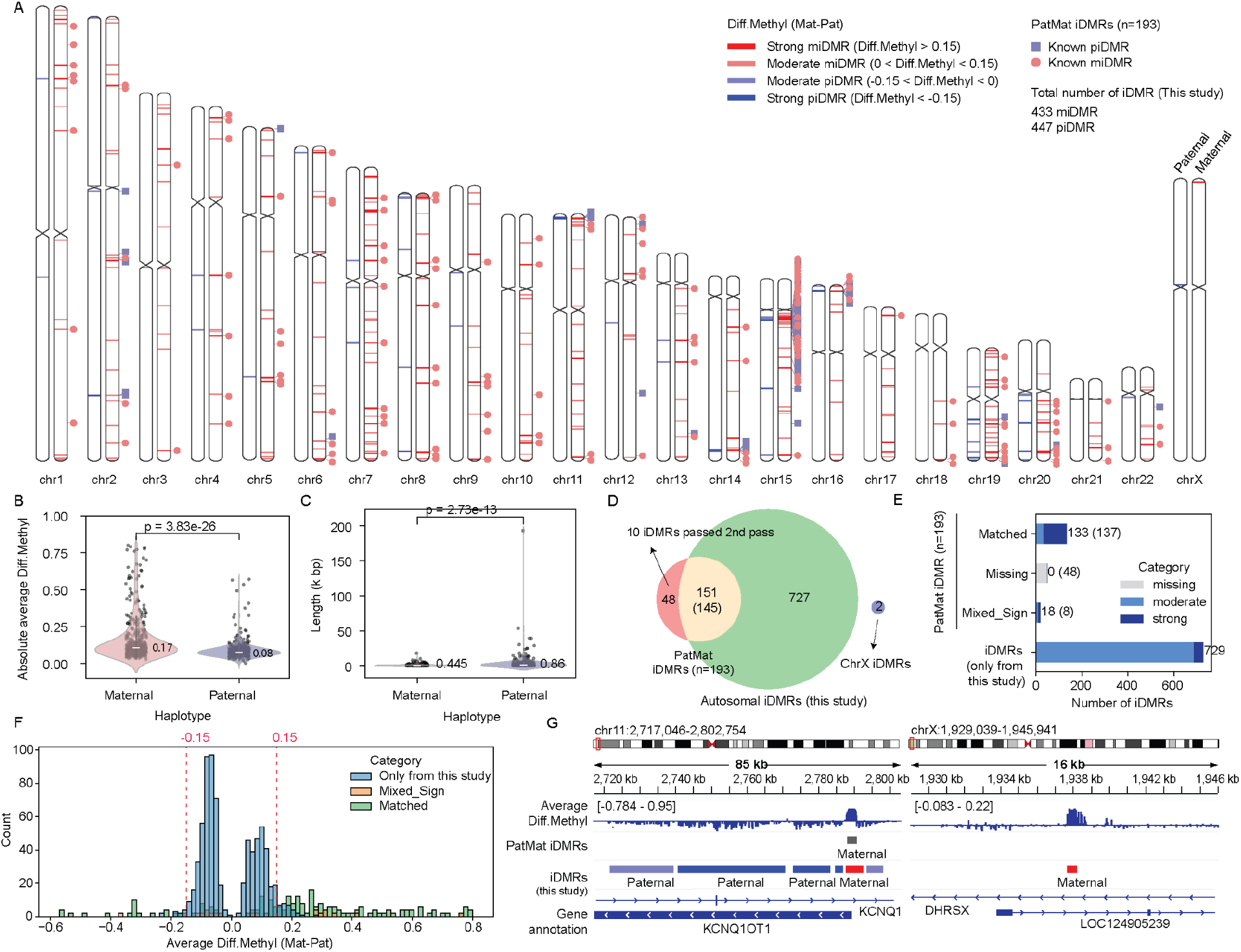
Categorization of iDMRs identified from 123 HPRC trios, including chromosome X. (A) Genomic overview of iDMRs identified in this study, colored by the magnitude and direction of differential methylation (maternal − paternal). PatMat iDMRs^19^ are shown alongside the chromosome ideograms as red circles (maternally hypermethylated) and blue squares (paternally hypermethylated). For visualization purposes, each iDMR was extended by 500 kb on both sides. (B) Absolute methylation differences and (C) length distributions of maternally and paternally hypermethylated iDMRs. P-values were calculated using the Wilcoxon rank-sum test. The difference remained significant after excluding the single outlier among piDMRs (P = 4.10 × 10⁻¹³), which overlaps a well-characterized ICR within the 15q11-13q locus. (D) Venn diagram showing the overlap between PatMat iDMRs and those identified in this study. iDMRs on chrX are shown separately. Numbers indicate iDMRs identified in this study, with unique PatMat iDMR counts in parentheses. Among the 48 PatMat iDMRs not recovered in the initial analysis, 10 exhibited significant methylation differences between parental haplotypes in region-based paired t-tests, with directions concordant with previous reports. (E) Number of iDMRs identified in this study and those overlapping the 193 PatMat iDMRs. Numbers indicate iDMRs identified in this study, with unique PatMat iDMR counts in parentheses. Colors indicate differential methylation categories (strong or moderate), with gray representing missing iDMRs. The y-axis shows overlap categories: “Matched” indicates overlapping iDMRs with concordant hypermethylated parental haplotypes; “Mixed-sign” indicates PatMat iDMRs overlapping multiple iDMRs identified in this study with opposing methylation directions; and “Missing” indicates PatMat iDMRs with no overlap. (F) Distribution of differential methylation with the thresholds used to distinguish strong versus moderate iDMRs indicated by red dotted lines (-0.15 and 0.15). (G) The “Average diff. methylation” track displays the mean methylation difference between maternal and paternal haplotypes (Mat − Pat) for each CpG. The “PatMat iDMRs” track shows regions identified in previous studies. The “iDMRs (This study)” track displays iDMRs identified in this study, colored according to their classification categories (as in Fig. 1A). The “Gene Annotation” track shows genes and transcripts based on the latest T2T-CHM13v2.0 gene annotation (JHU RefSeq + Liftoff v5.3). The left panel shows iDMRs within the *KCNQ1OT1* and *KCNQ1* loci, while the right panel shows iDMRs within the *DHRSX* and *LOC124905239* loci.

Comparison of iDMRs by haplotype showed that miDMRs exhibited larger absolute methylation differences than piDMRs (Fig. 1B), as reflected by a greater number of strong miDMRs identified in this study (Supplementary Fig. 3) and the predominance of maternally hypermethylated iDMRs in PatMat iDMR (145 maternal versus 48 paternal). In contrast, piDMRs tended to span longer genomic regions than miDMRs (Fig. 1C). The longest iDMR was paternally hypermethylated and located at 15q11.2, corresponding to a well-known imprinting control region (ICR) in the pericentromeric region of chr15 (Supplementary Fig. 4)^45^. The difference in length distributions remained significant after excluding this longest piDMR (P = 4.10 × 10⁻¹³; Supplementary Fig. 5).

To benchmark our *ab initio* iDMR discovery in HPRC samples, we compared our calls with a curated reference set of 193 iDMRs previously compiled from multiple studies and used for haplotype assignment by the “PatMat” method^19^. Our *ab initio* HPRC analysis recovered 145 of these PatMat iDMRs, including 137 with a concordant methylation direction and eight that showed mixed-sign methylation patterns (Fig. 1D,E). The PatMat iDMRs with mixed methylation patterns in the HPRC data appear to mark a true iDMR, but extend too far into the surrounding sequence and are broken into multiple regions by our analysis (Supplementary Fig. 6 and 7).

These differences in iDMR boundaries may reflect the greater sensitivity and finer resolution of our approach in delineating iDMR boundaries. Of the 48 PatMat iDMRs we did not recover, 46 (95.8%) were supported by only a single study (Supplementary Fig. 8A). Paired t-tests comparing the mean methylation levels of maternal and paternal haplotypes within each of the 48 unrecovered regions revealed significant differences in just 10 regions (Supplementary Fig. 8B). We saw no evidence of differential methylation in the previously reported direction at the remaining 38 regions, which could reflect biological differences between the cell types tested and/or false positives in the prior studies. Of the 10 significant regions, the two with the strongest statistical support (FDR < 0.001 for both) were also the only two of the 48 reported by multiple studies. Closer inspection revealed that both lay near telomeres, where individual CpGs lacked methylation measurements for one or more haplotypes. These gaps prevented recovery of either region because our *ab initio* analysis required methylation measurements for both haplotypes in every sample at each CpG tested. We obtained similar results when comparing our iDMRs with 229 iDMRs previously identified by deCODE using parent-of-origin-phased blood methylomes^14^, recovering approximately 80% of these regions (Supplementary Fig. 9).

Although our stringent coverage requirement may limit discovery in difficult-to-map regions near telomeres and centromeres, we recovered most iDMRs in both reference sets, strengthening confidence in our ab initio iDMR calls.

We identified 729 iDMRs not present in the previously reported PatMat set, of which 688 exhibited moderate differential methylation (Fig. 1E and 1F). Interestingly, several novel iDMRs clustered around PatMat iDMRs but showed the opposite direction of parent-specific methylation (Fig. 1G). We also identified candidate imprinted loci with strong iDMRs located within or near gene promoters (within 1 kb), including *GPR39*, *NPY*, and *ZNF890P* (Supplementary Fig. 10). Notably, *GPR39*, a zinc-responsive G protein-coupled receptor, has been reported as a downstream target of *PLAGL1*, a paternally expressed imprinted transcription factor^46,47^. Similarly, *NPY* encodes a neuropeptide involved in appetite regulation, stress responses, and metabolic homeostasis, and has been linked to regulatory pathways involving *ZDBF2*, a paternally expressed imprinted gene^48,49^. Furthermore, *ZNF890P* has previously been proposed as a candidate imprinted locus based on expression quantitative trait locus (eQTL) analysis^50^. We identified a total of 50 previously unreported genes with moderate iDMRs covering their promoters (Supplementary Table 4).

The availability of the HPRC’s fully phased diploid assemblies provided an opportunity to investigate parent-of-origin-specific methylation on chrX, where no well-established iDMRs have previously been reported. We identified two candidate iDMRs, including a strong miDMR within an intron of *DHRSX* near *LOC124905239* (Fig. 1A and 1G), and a moderate piDMR located near the centromere (Fig. 1A). Notably, the *DHRSX* locus has previously been identified as a putative ICR based on full methylation in oocytes, absence of methylation in sperm, and intermediate methylation (∼50%) in embryonic brain, kidney, and liver^45^.

### Pooch accurately predicts parent of origin for all chromosomes

We next trained an elastic net regression classifier to predict parent of origin for each chromosome pair using differential methylation in the HPRC dataset. To capture methylation differences between parental homologs, we present haplotypes to the classifier as pairs (Hap1, Hap2), with each CpG represented by the methylation difference Hap1 − Hap2. This paired representation draws on both maternally and paternally hypermethylated loci, enabling parent-of-origin assignment even for chromosomes with few paternally hypermethylated iDMRs, as most chromosomes contain loci with strong maternal hypermethylation. The classifier distinguishes between the two possible parental configurations, with Mat − Pat designated as the positive class (label = 1) and Pat − Mat as the negative class (label = 0). (Fig. 2A). To comprehensively capture parent-of-origin methylation signals, including subtle differences that might not be detected by DSS^44^, the classifier was trained on all individual CpGs present in at least 95% of training haplotypes rather than the iDMRs discussed above. All training data was presented in both haplotype orientations to balance the classes.

**Fig. 2:**
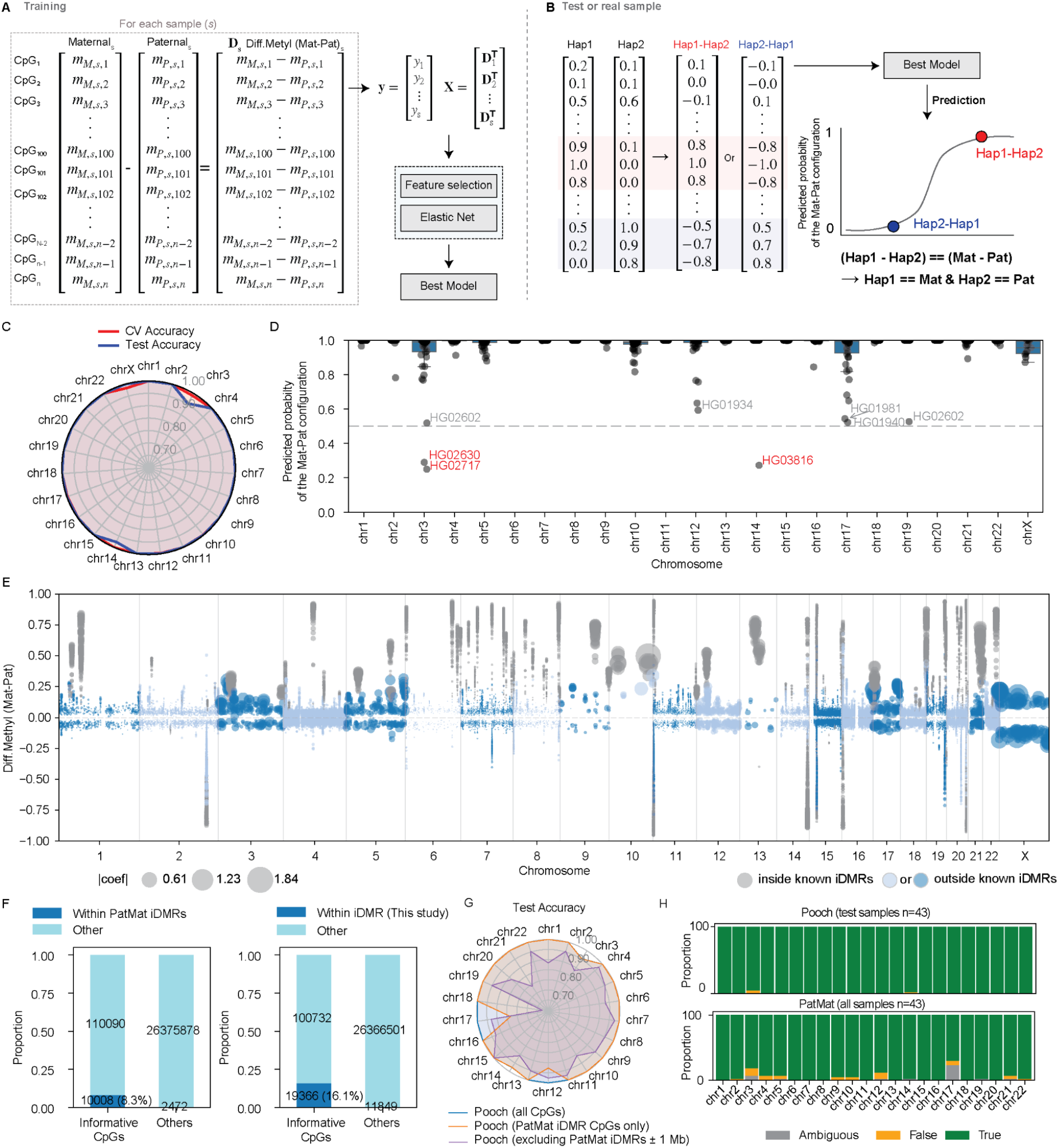
Pooch training and parent-of-origin prediction across training. (A) Model training. Diff.Methyl (D*_s_*) was calculated as both Mat − Pat for each CpG site, where *m* denotes methylation proportion, *M* and *P* represent maternal and paternal haplotypes, *s* denotes the sample in the training set (80 for autosomes and 46 for chromosome X), *n* is the number of CpGs per chromosome, and D*_s_*^T^ represents the transposed D*_s_* vector for sample *s*, allowing it to be paired with the corresponding haplotype configuration label *y*. (B) Model prediction. Diff.Methyl was calculated between haplotypes of unknown parental origin (Hap1 − Hap2 or Hap2 − Hap1) and used to predict the probability that the input configuration corresponds to Mat − Pat. (C) Accuracy was evaluated using nested cross-validation on the training dataset (red) and test accuracy (blue). (D) Predicted probability of the Mat − Pat haplotype configuration for all held-out test samples. Samples with probabilities below 0.6 are labeled in grey and those below 0.5 labeled in red. (E) Differential methylation (Diff.Methyl; Mat − Pat) across chromosomes. Point size indicates the absolute value of the learned model coefficient. (F) Left: Occurrence of model-learned informative CpGs within and outside of PatMat iDMRs. Right: Occurrence of informative CpGs within and outside of all iDMRs identified by this study. (G) Test accuracy of models trained using all CpGs, only CpGs within PatMat iDMRs, and only CpGs outside of PatMat iDMRs by at least 1 Mb. (H) Percentage of samples correctly and incorrectly classified by Pooch and the previously published “PatMat” method^19^.

We divided the 123 samples into 80 training samples and 43 held-out test samples for the autosomes (Supplementary Table 1, Supplementary Fig. 11), and 46 training samples for chromosome X, balancing those with higher maternal (n = 23) and paternal (n = 23) *XIST* promoter methylation to minimize XCI bias, while reserving the remaining 13 samples for testing (Supplementary Table 2 and Supplementary Fig. 12). We used the same elastic net classifier for the autosomes and chromosome X, with an additional preprocessing step for chromosome X to account for XCI. Specifically, we fitted a linear regression model at each CpG site to estimate the contribution of *XIST* activity (as measured by *XIST* promoter methylation) on the differential methylation of each ChrX CpG site. The estimated contribution was then subtracted from the observed methylation differences to obtain XCI-adjusted parent-of-origin methylation signals, which were subsequently used as input to the elastic net classifier. For both the autosomes and chromosome X, five-fold cross-validation (CV) was performed on the training samples, including hyperparameter optimization and independent feature selection within each fold to select differentially methylated CpGs using the ANOVA F statistic at a false discovery rate (FDR) of 0.05 (Supplementary Fig. 13). The final model for each chromosome was refitted on all training samples using the optimal hyperparameters and evaluated on the held-out test set.

For all held-out test samples, the haplotype configuration was consistently presented as Mat − Pat for ease of presentation (Fig. 2B), with equivalent results confirmed for the reverse orientation. The classifier achieved high accuracy during both validation and testing, with a mean nested CV accuracy of 0.999 and test-set mean accuracy of 0.997 (Fig. 2C, 2D and Supplementary Table 5), without any apparent population-specific performance degradation (Supplementary Fig. 14). Chr3 showed relatively lower test accuracy, but still achieved 95.23% (Fig. 2C and 2D). Notably, Pooch achieved perfect test accuracy for chrX and chr17, for which zero and one PatMat iDMR had previously been reported, respectively (Fig. 2C and 2D).

We investigated the samples with low prediction probabilities (p < 0.6) and false-negative classifications (p < 0.5) from otherwise well-performing chromosomes. The one false-negative sample for chr14, from HG03816, contained a long homozygous region with few parent-specific k-mers within the strongest imprinted methylation region (Supplementary Fig. 15A and 15B).

This resulted in ambiguous read alignments, even for the long ONT reads (Supplementary Fig. 16), preventing accurate methylation calling for the individual haplotypes. In another example, the HG02602 assembly contained large missing regions on chr19 (Supplementary Fig. 17), and its diploid graph transitions from a diploid structure to a single haplotype, suggesting a potential cell-line artifact (Supplementary Fig. 18). These examples illustrate that while Pooch achieves high accuracy, it relies on the underlying haplotypes to be correctly assembled and the ONT data to be confidently mapped to the correct haplotype, which will depend on both read length and assembly accuracy.

We hereafter refer to CpGs with nonzero coefficients after elastic net regularization as “informative” CpGs (Supplementary Fig. 13), which contribute to prediction in our fitted model but do not necessarily encompass all CpGs carrying parent-of-origin methylation signals.

Comparison of their genomic distribution with that of the PatMat iDMRs revealed that Informative CpGs with large model absolute coefficients frequently overlapped these regions, with more large positive coefficients than negative ones, consistent with a greater number of strong miDMRs than piDMRs (Fig. 2E). Informative CpGs were also significantly enriched within iDMRs (p < 0.001, Supplementary Fig. 19), and the proportion of informative CpGs overlapping all iDMRs identified in this study was 16.1%, approximately twice the proportion overlapping the PatMat iDMRs (8.3%). However, the majority of informative CpGs (83.9%) remained outside these regions, distributed across the chromosomes (Fig. 2F). These results suggest that, although PatMat iDMRs are strongly enriched for predictive CpGs, other CpGs outside the annotated iDMRs also harbor parent-of-origin information.

To further evaluate the contributions of CpGs outside of PatMat iDMRs, we trained and tested five additional models, one using only CpGs within PatMat iDMRs and four excluding CpGs within 1 bp, 2 kb, 100 kb, or 1 Mb of PatMat iDMRS. As expected, the model restricted to PatMat iDMRs performed the best overall with mean accuracy 0.986, and the exclusion models retained predictive accuracy above 0.92, supporting the existence of parent-of-origin methylation signals beyond the boundaries of PatMat iDMRs (Fig 2G, Supplementary Fig. 20-24, and Supplementary Table 6 and 7). As progressively larger regions surrounding PatMat iDMRs were excluded, the models placed greater weight on CpGs in regions with moderate differential methylation that had small (but nonzero) coefficients in the model trained using all CpGs (Supplementary Fig. 25). Such regions were found scattered throughout the chromosome body. In the case of the model restricted to only PatMat iDMRs, accuracy was decreased for chr12 and chr17, suggesting important regions for parent-of-origin classification are missed by models that focus only on the PatMat iDMRs.

We finally compared the performance of Pooch to PatMat, a previously described strategy that assigns haplotypes based on methylation levels at CpGs within autosomal iDMRs^19^. Across the 945 autosomal scaffolds from 43 test samples, PatMat misassigned 25 (2.64%) and produced ambiguous assignments for 17 (1.79%), achieving an average accuracy of 95.56, whereas Pooch achieved a test accuracy 99.68 (Fig. 2H and Supplementary Table 8). Chr17 showed reduced assignment accuracy with PatMat, resulting in 6.97% false-negative and 23.25% ambiguous assignments. In contrast, Pooch correctly assigned the haplotypes of all chr17 scaffolds, owing it its expanded feature set. PatMat also failed to correctly classify two of the three scaffolds that Pooch misclassified (Supplementary Fig. 26). In contrast, Pooch correctly classified 40 pairs for which PatMat produced incorrect or ambiguous assignments. Together, these results demonstrate that Pooch provides more robust haplotype assignment than PatMat, including for chrX and autosomes with a limited number of iDMRs.

### Pooch generalizes to different assemblies, sequencing platforms, and karyotypes

We next tested Pooch on a T2T pedigree using both HiFi sequencing from blood and ONT sequencing from LCLs for four members of an African American family including the grandmother (PAN010), grandfather (PAN011), their daughter (PAN027), and granddaughter (PAN028) (Supplementary Table 9)^51^. Pooch produced consistent parent-of-origin assignments across both ONT LCL and HiFi blood datasets, with only one discrepancy between the two sequencing platforms (chr9 from PAN010). All predictions further agreed with the prior study’s haplotype assignments except chr17 in PAN011 (Fig. 3A, Supplementary Table 10), where Pooch predicted a Hap1 maternal-origin probability of only 0.01 and 0.05 from ONT and PacBio, respectively. Unfortunately, since this is the grandfather, no parental markers are available for confirmation and the original study made its assignment by manual inspection of a single known iDMR. The Pooch results for both blood and LCL strongly suggest that the haplotype labels for this chromosome should be swapped.

**Fig. 3:**
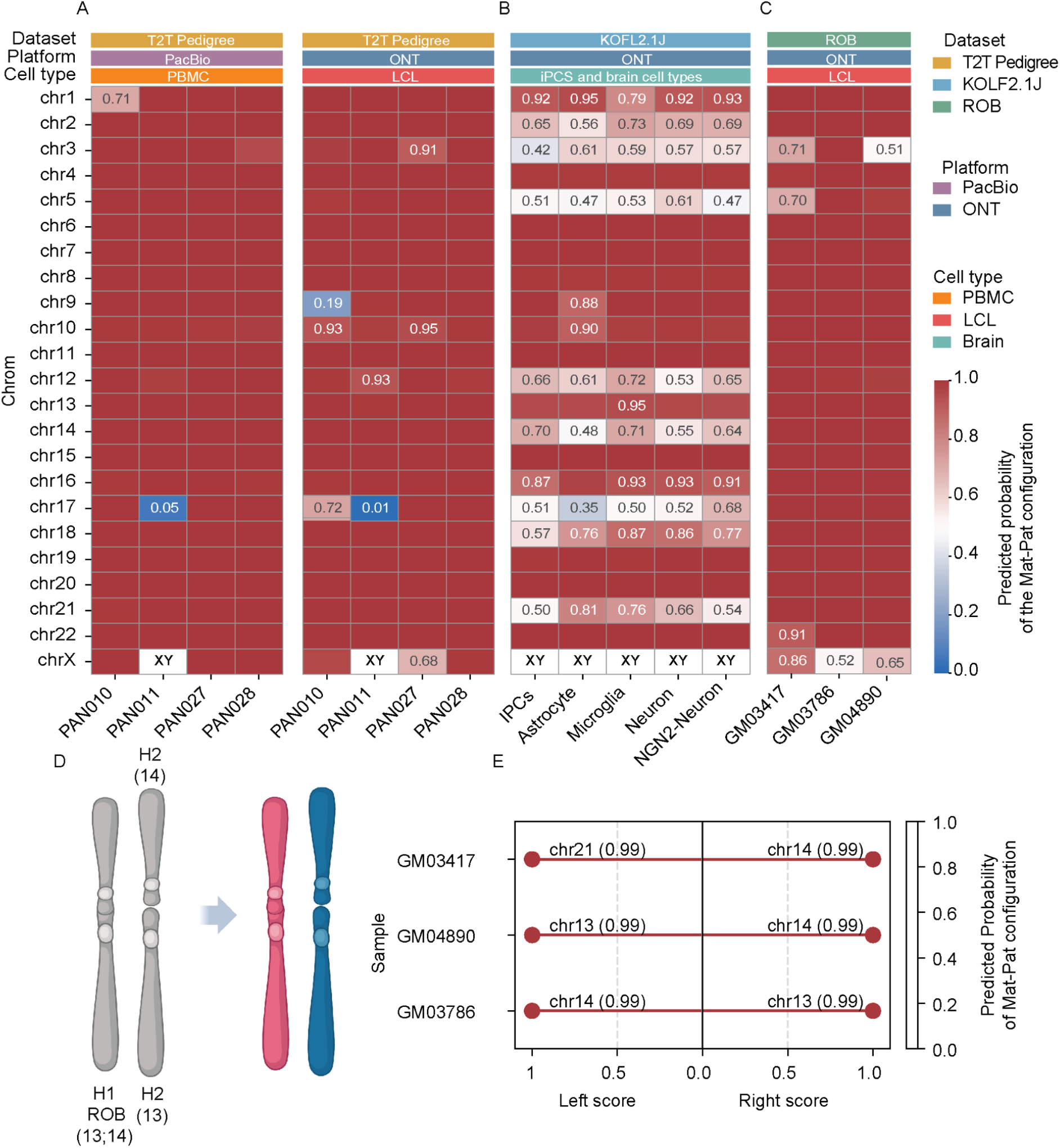
Applying Pooch on abnormal karyotype samples. (A) Pooch prediction results for four samples from a T2T pedigree. (B) Pooch prediction results for the KOLF2.1J dataset, comprising multiple cell types, including induced pluripotent stem (iPS) cells, cortical neurons, NGN2-induced neurons (NGN2), astrocytes and microglia. (C) Pooch prediction results for Robertsonian translocation (ROB) samples. (A-C) Colors indicate concordance with a Mat − Pat configuration, with values below 0.95 highlighted and annotated with their corresponding numerical values. (D) Schematic diagram illustrating the application of Pooch to ROB chromosomes. (E) Pooch predictions for the ROB chromosome arms. The left and right scores indicate the Pooch prediction for the two long arms that form each ROB chromosome, in the order shown.

Methylation levels are context-dependent and can vary across cell types and cell-cycle stages, so we also tested Pooch on KOLF2.1J induced pluripotent stem cells (iPSCs), for which a T2T assembly is available, as well as methylation data from derived brain cell types^52^. Because parental sequencing data were unavailable to establish the true parent-of-origin haplotypes, we used differential methylation between the primary and alternative haplotypes (pri − alt) as the input and assigned parental origin based on a majority vote of the Pooch predictions across cell types. Most chromosomes showed high-confidence predictions, with predicted maternal probabilities approaching p=1 (Fig. 3B). However, several chromosomes showed more ambiguous probabilities, including chr5, chr14, and chr21, potentially reflecting cell type-specific methylation differences or epigenetic changes associated with cellular reprogramming^53,54^.

Nevertheless, the predicted haplotype direction was largely consistent across cell types, with the exception of chr3 and chr17, as before. Overall, Pooch showed the highest confidence in the LCL cell type used for training, but still produced consistent classifications in other cell types, including reprogrammed iPSCs and differentiated brain cells, albeit with increased uncertainty for some chromosomes.

Parent-of-origin methylation patterns associated with genomic imprinting are highly conserved and are generally thought to be maintained despite chromosomal copy-number abnormalities, such as monosomy or uniparental inheritance^55^. However, we sought to determine whether these patterns are preserved in genomes with abnormal karyotypes, such as Robertsonian translocations (ROBs). To investigate, we analyzed previously published T2T ROB assemblies generated without trio data and their corresponding ONT sequencing data from fibroblast cell lines^36^ (Supplementary Table 11 and Supplementary Fig. 27). Methylation profiles of the fusion chromosomes were lifted over to their corresponding chromosomes in the T2T-CHM13v2.0 reference (Fig. 3D, Supplementary Fig. 27), generating chromosome-specific methylation profiles for each half of the fused chromosomes. We then applied Pooch to determine whether the two components of each fusion chromosome could be consistently assigned to the same parental haplotype and to infer their parental origin. Pooch assigned the haplotype configuration with high confidence and predicted a maternal-origin (p=0.99) for all components of the ROB fusions (Fig. 3C and 3E), consistent with a higher frequency of Robertsonian translocations arising during oogenesis^56^. Together, these results demonstrate that Pooch can determine parent of origin for multiple assemblies, sequencing technologies, cell types, and karyotypes.

### Imprinting patterns on chr3, chr17, and chrX

The prior iDMR annotation from PatMat included an average of 8.7 iDMRs per chromosome, with zero, one, and two iDMRs on chrX, chr17, and chr3, respectively. Despite the lack of previously annotated iDMRs, Pooch achieved high parent-of-origin classification accuracy for all three chromosomes (Fig. 2C and 2D), suggesting that it had recognized previously unreported imprinting signals. We therefore examined the genomic distribution of our DSS-identified iDMRs and Pooch model coefficients for these specific chromosomes. On chr3, we identified three strong miDMRs and 11 moderate iDMRs, of which six overlapped clusters of strong model coefficients (Fig. 4A). Two of these regions overlapped a PatMat iDMR near the *MOBP* and *FGF2* gene (Supplementary Fig. 28), and we further identified candidate iDMRs and strong coefficient clusters near the promoters of *SSUH2*, *GADL1*, and *COL7A1* (Fig 4A). *GADL1* has been reported as a rat-specific imprinted locus with paternal allele-specific expression, although imprinting has not been established in humans^57^, whereas *SSUH2* and *COL7A1* were not previously implicated in genomic imprinting. On chr17, we identified four distinct peaks of model coefficients that overlapped with our candidate iDMRs, one of which colocalized with a PatMat iDMR near the *CHRNE* gene^14^ (Supplementary Fig. 28). Two additional peaks were located near the promoter of *LIAT1* and *TNRC6C*, respectively, representing previously unreported candidate iDMRs (Fig. 4A).

**Fig. 4:**
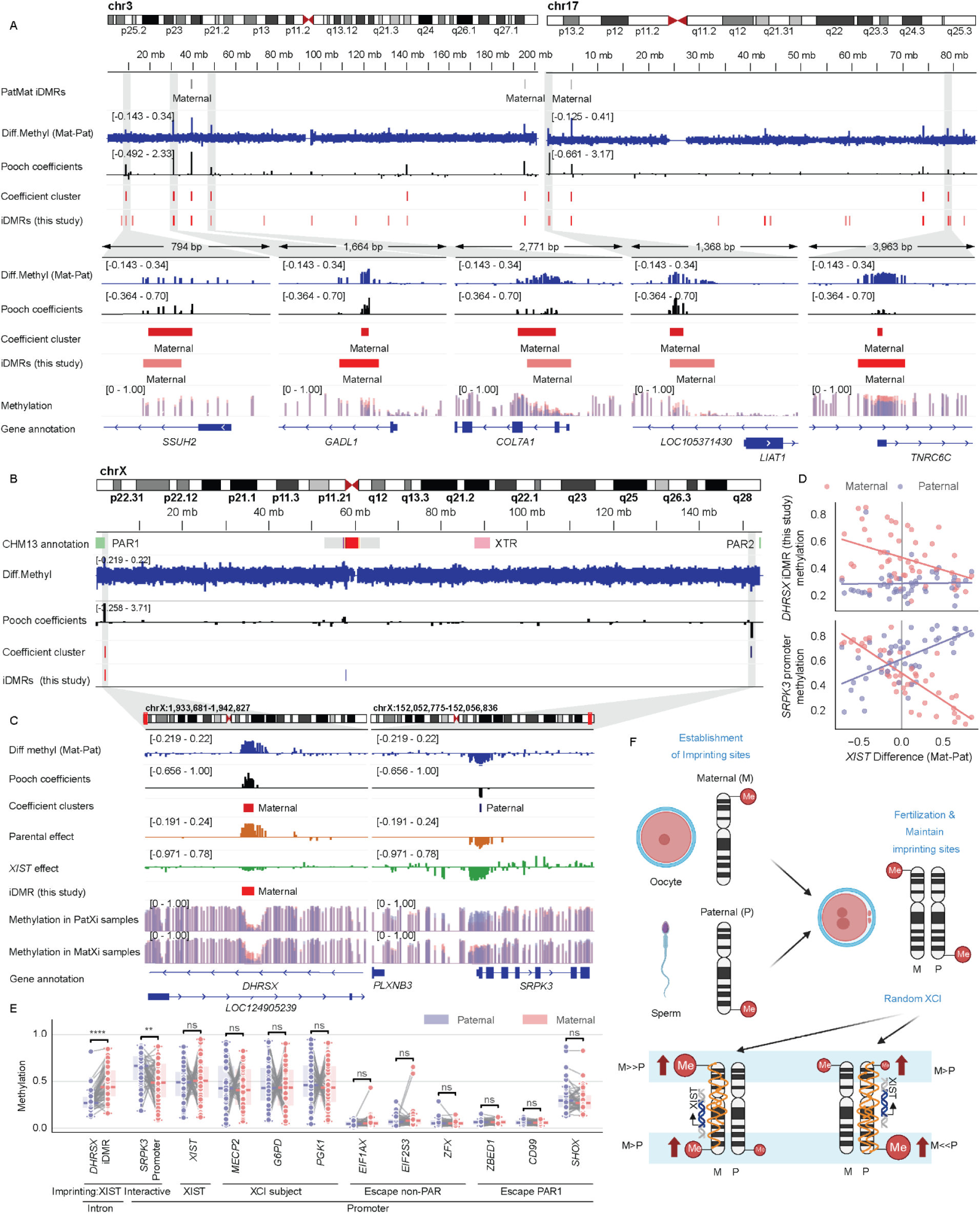
Detailed characterization of imprinting on chr3, chr17 and chrX. (A) Genomic overviews of chr3 and chr17, with zoomed-in views of iDMRs overlapping promoter regions shown below with tracks for average differential methylation “Diff.Methyl”, windowed sums of Pooch model coefficients, and DSS-identified iDMRs. Model coefficients were summed within non-overlapping 100-bp bins for visualization, and the clusters highlight regions containing coefficients of the same sign separated by no more than 500 bp and containing at least 3 CpGs. Methylation tracks show overlaid bar charts of maternal (red) and paternal (blue) methylation fractions. (B) Genomic overview of chromosome X with the same tracks as panel A. (C) Zoomed-in genomic view of the *DHRSX* intron (left) and *SRPK3* promoter (right) containing a cluster of large model coefficients. The “Parental effect” and “*XIST* effect” tracks show the estimated contributions of each term to CpG methylation differences during model training. The “Pat Xi” and “Mat Xi” tracks show the mean methylation levels of maternal (pink) and paternal (blue) haplotypes in samples with lower paternal and maternal *XIST* promoter methylation, respectively. (D) Haplotype-specific methylation levels for individual samples plotted against *XIST* promoter differential methylation, with points colored by parental haplotype. Note, greater maternal Xi is a difference < 0 and greater paternal Xi is a difference > 0. (E) Paired t-test comparisons of maternal and paternal methylation levels at the *DHRSX* iDMR and selected gene promoters, including *SRPK3*, *XIST*, and genes known to undergo or escape XCI. (F) Schematic illustration of the relationship between genomic imprinting and random XCI on the human X chromosome, whereby the observed methylation of *DHRSX* and *SRPK3* depend both on the parent of origin and XCI.

In humans, XCI inactivates all but one X chromosome through the activity *XIST*, so cell or tissue-specific inactivation can confound any parent-of-origin effects. As noted above, some HPRC LCLs appear to exhibit clonally skewed XCI, which can be measured and corrected for by examination of the *XIST* promoter. Using the corresponding Pooch model, we identified two clusters of informative coefficients on chrX, one of which exhibited higher maternal methylation (positive coefficients) within an intron of the *DHRSX* gene (Fig. 4B and 4C, Supplementary Fig. 29 and 30), and the other exhibited higher paternal methylation (negative coefficients) near the *SRPK3* promoter (Fig. 4B and 4C). Interestingly, both regions exhibited XCI-dependent methylation patterns.

To distinguish parent-of-origin effects from XCI-associated methylation, we performed an interaction analysis incorporating parental haplotype, *XIST* promoter methylation, and their interaction. We first analyzed 94 of the 95 regions on chromosome X previously proposed as candidate ICRs^45^, excluding one region that could not be lifted over to T2T-CHM13v2.0 coordinates (Supplementary Fig. 31). A total of four regions exhibited significant haplotype effects (FDR < 0.05), among which the region overlapping the *DHRSX* iDMR showed the strongest association with parental haplotype (FDR = 1.79E-12; Supplementary Table 12 and Supplementary Fig. 31). Although the direct effect of *XIST* promoter methylation was not significant for *DHRSX* (FDR = 0.59), the haplotype-by-*XIST* interaction was significant (FDR =0.001), and *DHRSX* showed increasing maternal methylation with maternal X inactivation but consistently low methylation on the paternal haplotype regardless of XCI status (Fig. 4D). This difference may reflect the location of *DHRSX* within PAR1, which generally escapes XCI. In contrast, the *SRPK3* promoter exhibited significant effects of parental haplotype (FDR = 3.38E-04), *XIST* promoter methylation (FDR = 6.88E-09), and their interaction (FDR= 2.71E-28) (Supplementary Table 12 and Supplementary Fig. 31). Unlike the *DHRSX* iDMR, the *SRPK3* promoter largely tracked XCI status, with increased methylation observed on the inactivated chromosome, suggesting a greater influence of XCI-associated methylation (Fig. 4D and Supplementary Fig. 32). Paired t-tests further revealed significant methylation differences between parental haplotypes at both the *DHRSX* iDMR and *SRPK3* promoter, while all other loci followed the expected pattern of random XCI (Fig. 4E). Extending our analysis beyond putative ICRs to all CpGs across chrX also recovered prominent association signals for *DHRSX* and *SRPK3* (Supplementary Fig. 33). Together with previously reported differential methylation patterns in gametes at these loci^45^ (Supplementary Fig. 34), our findings suggest that chromosome X harbors parent-of-origin-associated methylation signals in somatic tissues, whose magnitude and direction is partially modulated by XCI (Fig. 4F).

Given that *DHRSX* is located within PAR1, which is shared between chrX and chrY, we extended our differential methylation analysis to XY samples. We compared DNA methylation between the maternally inherited X chromosome and the paternally inherited Y chromosome, with Y chromosome coordinates lifted over to chromosome X. Consistent with the maternal hypermethylation observed at the *DHRSX* locus in XX samples, XY samples also exhibited significantly higher methylation on the maternal X chromosome than on the paternal Y chromosome (paired t-test, P < 0.05), although the magnitude of the difference was substantially smaller than in XX samples (mean difference = 0.0881, Supplementary Fig. 35). The XY samples also showed overall higher levels of methylation on both haplotypes compared to the XX samples. Thus, methylation of this *DHRSX* locus appears to also be affected by sex-specific effects, but with our *XIST*-based correction method, it remains a highly informative site for parent-of-origin prediction.

To assess whether the interactions between parent of origin and XCI described above affected predictive performance, we compared the XCI-adjusted and unadjusted models using T2T pedigree female samples (n = 6) and the HPRC held-out test set (n = 13). The T2T pedigree samples had ONT and PacBio data derived from different tissues of the same individuals, with distinct XIST promoter methylation patterns (Supplementary Fig. 36A). While both models correctly classified the X chromosome haplotypes, the XCI-adjusted model assigned higher probabilities to the correct parental configuration than the unadjusted model (Supplementary Fig. 36B and 36C, Supplementary Table 13), highlighting the importance of accounting for *XIST* promoter differential methylation for reliable chrX parent-of-origin assignment.

### Pipelines and computational benchmark

For ease of application, we developed a Snakemake pipeline that integrates all steps of the Pooch workflow, including inference using the HPRC-trained model described here as well as training for new samples and species (Fig. 5A). For human samples, the Pooch pipeline supports four input types for methylation analysis: (i) raw FASTQ files, (ii) BAM files aligned to a personalized genome, (iii) haplotype-specific BED files based on personalized-genome coordinates, and (iv) haplotype-specific BED files based on T2T-CHM13 coordinates (Fig. 5A). The pipeline automatically recognizes the input format and starts from the corresponding step of the workflow. A chromosome-scale, phased diploid assembly is required for each sample to determine haplotype-specific methylation calls.

**Fig. 5:**
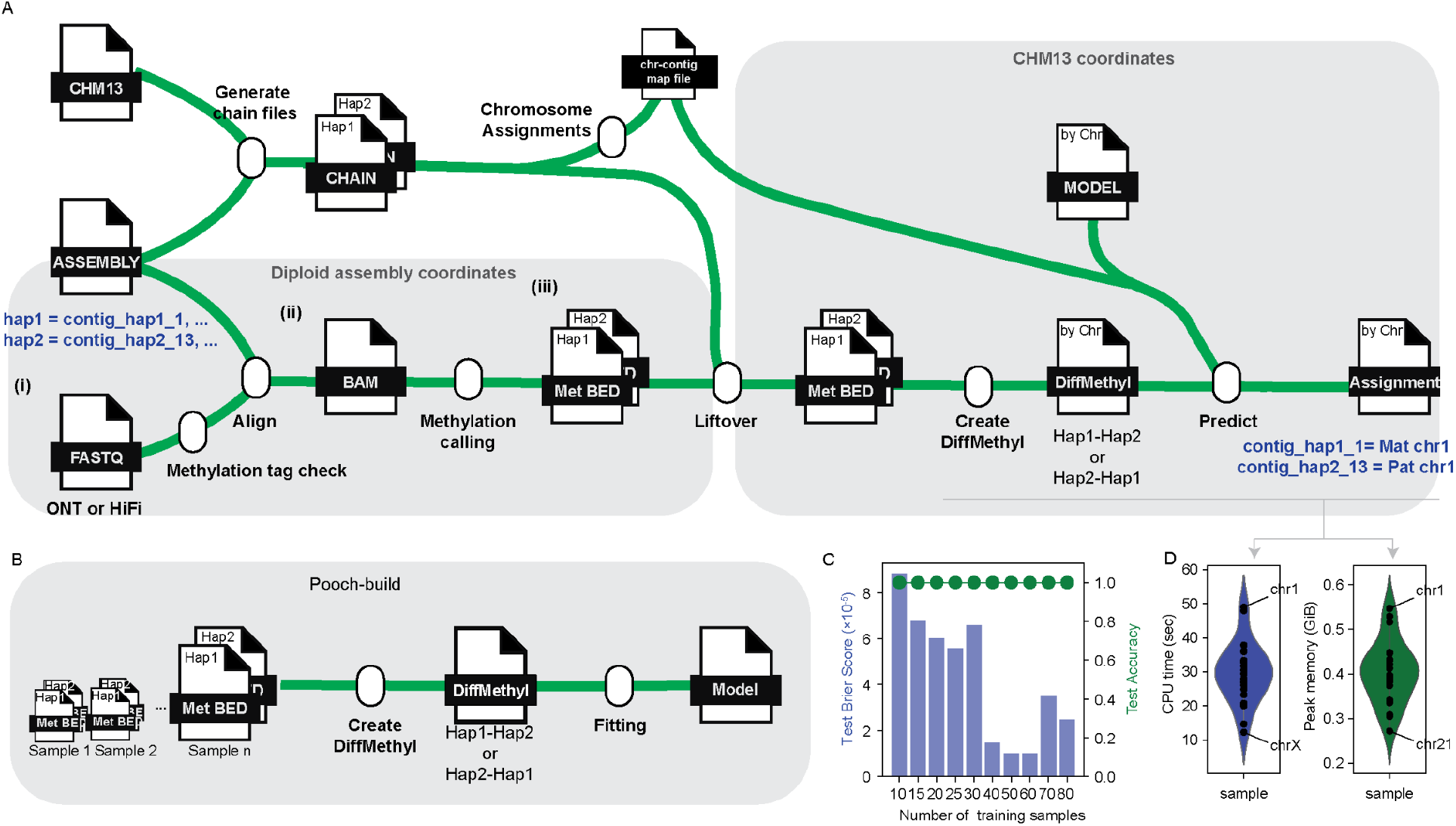
Pooch Snakemake pipeline and computational resource requirements. (A) End-to-end Pooch prediction pipeline with computation steps as nodes and inputs/outputs as documents on the edges. (B) Pooch-build pipeline for training customized Pooch models. (C) Prediction performance as a function of the number of training samples using chr22 as a test case. Model performance was evaluated using test accuracy and test Brier score (mean squared error, lower is better). (D) Computational resources required for the Pooch prediction step, including CPU time and memory usage for each chromosome. Chromosomes with the highest and lowest values for each metric are highlighted.

For raw sequencing data, reads containing methylation tags are first aligned to a personalized diploid genome, followed by methylation calling at CpG sites (Fig. 5A). Haplotype-specific chain files are then generated between each personalized genome and T2T-CHM13v2.0. These chain files are used to assign each scaffold to its corresponding chromosome, generating a chromosome-contig mapping file that is automatically used during the prediction step. CpG methylation calls are subsequently lifted over to T2T-CHM13v2.0 coordinates using the chain files, resulting in haplotype-specific methylation profiles for Hap1 and Hap2.

Because genomic imprinting is observed across cell types and species, Pooch models can be retrained for other cell types and organisms. To facilitate model customization, we developed Pooch-build, a training pipeline for generating custom Pooch models (Fig. 5B). Notably, for chr22, models trained with as few as 10 samples achieved perfect accuracy, demonstrating that Pooch-build can efficiently generate high-performing models with relatively small training datasets (Fig. 5C).

Computational resource requirements vary depending on sequencing coverage and input data format. To assess computational performance, we benchmarked Pooch runtimes using ONT methylation data from HG002 for autosomal chromosomes and HG01081 for chromosome X (since HG002 is male). The most computationally demanding stage was preprocessing, which included read alignment and chain-file generation. In contrast, Pooch prediction was computationally efficient, requiring an average of 29.64 CPU-seconds per chromosome and approximately 0.40 GiB of peak memory on an Intel Xeon Platinum 8580 processor (Fig. 5D, Supplementary Table 14). These results demonstrate the computational efficiency of Pooch for parent-of-origin haplotype assignment without trio data and highlight its potential as a general framework for identifying methylation-based signatures of genomic imprinting beyond previously characterized iDMRs.

## Discussion

Genomic imprinting is a well-established epigenetic phenomenon that has previously been used to determine the parent of origin for haplotype-resolved assemblies^19^. However, only a few iDMRs have been previously reported on certain autosomes, and none have been well established on chromosome X, limiting the accuracy of prior approaches. In this study, we generated *ab initio* a comprehensive catalog of iDMRs using a collection of trio-based assemblies from the HPRC, recovering 145 of 193 PatMat iDMRs and identifying 729 previously unreported iDMRs, including 727 on autosomes and two on chrX. Most are of moderate differential effect, consistent with a continuous rather than binary model of imprinting^14^.

We leveraged these genome-wide methylation differences to develop Pooch, a classifier for determining the parent of origin for assembled haplotypes without requiring trio sequencing. Pooch achieved a mean test accuracy of 0.997, with accuracies exceeding 0.95 even for chr3, chr17, and chrX, which contain few strong iDMRs. Notably, 91% of model-informative CpGs were located outside PatMat iDMRs, and models trained exclusively on CpGs well outside of these previously reported iDMRs retained accuracy above 0.9, supporting the existence of parent-of-origin-associated methylation patterns beyond the strongly imprinted regions.

Differential methylation on chromosome X was substantially weaker than the autosomes, likely reflecting the influence of random XCI. Nevertheless, we identified two prominent regions of parent-of-origin-associated methylation within 2 Mb of each end of chrX. The strongest signal was detected within an intron of *DHRSX* in PAR1, which showed higher methylation in maternal haplotypes, especially for the inactivated maternal X. In contrast, slightly proximal to PAR2, the *SRPK3* promoter exhibited both paternal and XCI-associated methylation effects, with the direction and magnitude of differential methylation varying according to XCI status. Although both regions were influenced by XCI, the *DHRSX* locus appeared less affected, potentially reflecting its location within PAR1, a region containing numerous genes that escape XCI. Together, these findings suggest that parent-of-origin-associated methylation signals persist on chromosome X but are obscured by random XCI, which can be corrected for the purposes of classification.

Despite its broad applicability, Pooch has a few limitations. First, long stretches of highly homologous sequences may lead to ambiguous read mapping, even in fully resolved genome assemblies, potentially affecting parent-of-origin predictions when these regions overlap CpGs with large model coefficients. Second, Pooch’s dependence on coordinate liftover between personalized genomes and the T2T-CHM13v2.0 reference excluded the detection of possible iDMRs in complex genomic regions including centromeres, segmental duplications, structurally variable, and subtelomeric regions.

Pooch models were specifically trained using LCL-derived datasets from the HPRC, with the intent of classifying additional pangenome samples that lack a trio. Despite being trained exclusively on LCL-derived data, the model made consistent parent-of-origin predictions for other cell types and tissues, including blood, fibroblasts, as well as multiple iPSC-derived brain cell types. The model also performed well with both ONT and PacBio HiFi sequencing data.

These results highlight the robustness of Pooch beyond its primary target cell type and suggest its potential utility across diverse biological samples and sequencing technologies. Expanding the training datasets to encompass a broader range of cell types and tissues (e.g. peripheral blood) may further improve the generalizability of the model for clinical application.

Genomic imprinting is not restricted to humans but conserved across therian mammals and occurring in diverse taxa, including plants and insects^2^. To facilitate the application of Pooch across different species and datasets, we developed a pipeline for training custom models and developed an end-to-end Snakemake pipeline, including long-read alignment to diploid genome assemblies, model training, and parent-of-origin prediction. Once trained, Pooch classification is lightweight, facilitating its application to large numbers of samples including pangenome projects and T2T clinical genomes. Thus, it is now possible to generate complete diploid genomes, with maternal and paternal haplotype assignments, using a simple combination of long-read and Hi-C sequencing and without the need for trios.

## Methods

### HPRC assemblies and sequencing data download

We used diploid assemblies from 123 individuals in HPRC Release 2 for whom parental sequencing data were available^58^. The HPRC generated these assemblies with Verkko v2.2.1^59^, using trio markers to phase haplotypes and Hi-C data to scaffold the assemblies. The parent of origin assignments for these assemblies, as determined by the trio markers, were taken as the truth. We obtained the corresponding ONT sequencing data and sample metadata from the Human Pangenome Data Explorer (https://data.humanpangenome.org/raw-sequencing-data).

During Pooch validation, we identified reversed parental haplotype labels in three samples in the held-out test set (HG03579, HG02717, and HG02451). We traced these errors to the HPRC assembly metadata, which had interchanged the maternal and paternal identifiers. We corrected the labels in our analysis and used the corrected assignments throughout the study, including for model validation and iDMR calling.

We also identified a manual contig-splitting error during HPRC post-processing of the HG03942 assembly. The HPRC split the paternal contig pat-0000815 into chr12_pat-0000815_partA and chr19_pat-0000815_partB, but retained only 1 bp of the chromosome 12 segment in the assembly we analyzed. We therefore excluded chromosome 12 of HG03942 from all analyses because paternal reads could map to the maternal homolog and bias haplotype-specific methylation estimates.

After we reported these errors, the HPRC corrected the parental haplotype labels for HG02717 and the contig-splitting error in HG03942, as documented in the HPRC release notes (https://s3-us-west-2.amazonaws.com/human-pangenomics/submissions/6807247E-4F71-45D8-AECE-9E5813BA1D9F--verkko-v2.2.1-release2_asms/README.txt). When the HPRC submitted the higher-quality assembly for each individual to NCBI BioProject PRJNA1435250, it included the corrected assemblies for both samples. For HG03579 and HG02451, the HPRC submitted the higher-quality Hi-C-only assemblies, which required no correction. We archived all “trio-hic” assembled Verkko assemblies used for this study in the HPRC S3 bucket (https://s3-us-west-2.amazonaws.com/human-pangenomics/index.html?prefix=publications/Poo ch_2026/verkko-thic_assemblies). These archived files retain the original labels and contig sequences.

### Chain file generation and filtering

The personalized diploid assembly was first split by haplotype, and then chain files were generated by haplotype to T2T-CHM13v2.0 using a nextflow-based alignment and chain-building pipeline. The pipeline was executed via nextflow (nf-LO pipeline)^60^ with minimap2. Minimap2 alignment outputs were processed to generate a chainNet object via the nf-LO pipeline. The resulting chain file was segmented into smaller blocks using custom Python scripts, then converted to PAF to facilitate downstream processing.

PAF alignments were filtered and refined using rustybam utilities, as described previously^61^. segments were broken wherever indel lengths within them exceeded 10kb (break-paf), ends trimmed for quality (trim-paf -r) and inverted to improve alignment consistency (applied iteratively to alternate between forward and inverted orientations). The processed PAF was then converted back to chain format using paf2chain. The final chain file was inverted using custom Python scripts to produce a reciprocal mapping (target-to-source), enabling flexible coordinate liftover in both directions.

### Methylation calling and liftover to T2T-CHM13v2.0 for HPRC train and test set data

ONT reads were aligned to a personalized diploid genome assembly using Winnowmap^62^ (v2.03) with the parameters -x map-ont -I 8G --MD --cs -y -a -Y -L --eqx. FASTQ files were aligned in split batches and subsequently merged using SAMtools^63^ (v1.21). Primary alignments were retained by filtering with SAMtools using the -F 0×904 flag to exclude secondary, supplementary, and unmapped reads. Per-locus methylation levels and coverage were computed using modkit (v0.4.1) (https://github.com/nanoporetech/modkit), restricting the analysis to CpG sites with --preset traditional option set, the base modifications for the (+) strand C and the (-) strand C in CpG motifs were combined together on to the (+) strand position (see the modkit documentation for details) which causes the (-) strand positions to be offset by one, --cpg --combine-strands --ignore h.

Methylation levels derived from the personalized assembly were then lifted over to T2T-CHM13v2.0 coordinates using CrossMap^64^ (v0.7.0), with the haplotype-specific chain files described above. CpG liftover requires an additional coordinate adjustment because CpG sites are symmetric between the forward and reverse strands. When a CpG on the reverse strand is lifted over to the forward strand of the target reference (T2T-CHM13v2.0), the methylation call is mapped to the guanine (G) rather than the cytosine (C), resulting in a +1 bp coordinate shift. To correct for this, the chain files were separated into positive- and negative-strand alignment blocks and target coordinates for CpGs within negative-strand alignment blocks were shifted by -1 bp after liftover to restore the correct cytosine position.

### Classification of samples by the parental origin of the active X chromosome

To classify samples according to the parental origin of the Xi, we extracted methylated read counts and total coverage at CpG sites within the XIST promoter region (chrX:72285969 -72286319) for each haplotype across 59 female samples. For each haplotype, the coverage-weighted methylation proportion was calculated by dividing the sum of methylated read counts by the sum of total coverage across all CpG sites within the region. XIST differential methylation was then calculated by subtracting the paternal methylation proportion from the maternal methylation proportion (maternal − paternal). Samples with positive XIST differential methylation values were classified as having a paternally derived inactive X chromosome (pat Xi), whereas those with negative values were classified as having a maternally derived inactive X chromosome (mat Xi). No samples exhibited an XIST differential methylation value of zero.

### iDMR calling

We performed chromosome-specific differential methylation analysis using the DSS framework^44^ (v2.58.0) in R (v4.5.3). Methylated read counts and total coverage at individual CpG sites were extracted separately for maternal and paternal haplotypes across 123 samples for autosomal analysis and 50 samples for chromosome X analysis. For chromosome X, we selected a balanced subset comprising 25 samples with a mat Xi and 25 samples with a pat Xi, and combined into a BSseq object. CpG sites with missing methylation data in more than 5% of samples were excluded from further analysis.

To account for inter-sample variability and potential batch effects, we employed a paired design with sample identity included as a fixed effect. A design matrix incorporating haplotype and sample information was constructed, and the model was fitted using ‘DMLfit.multiFactor()’ with the formula ‘∼ haplotype + samplè. Differentially methylated loci (DMLs) were identified using ‘DMLtest.multiFactor()’ with the haplotype term, which evaluates haplotype-associated methylation differences using a Wald test. iDMRs were subsequently identified using ‘callDMR()’ with default parameters and a p-value threshold of 0.001.

To consolidate adjacent iDMRs potentially representing the same imprinted region, we merged iDMRs separated by no more than 1,000 bp that exhibited the same direction of haplotype-specific methylation differences. This post-processing step reduced the total number of iDMRs from 1,509 to 880.

### Comparison with PatMat iDMRs

We compared the set of 193 previously reported iDMRs^19^ (PatMat iDMRs) with the iDMRs identified in our analysis using BEDTools intersect in T2T-CHM13v2.0 coordinates. Identified iDMRs overlapping PatMat iDMRs were classified as matched if their parent-of-origin methylation directions were concordant and as mismatched if their directions were opposite. Identified iDMRs overlapping two or more PatMat iDMRs with discordant parent-of-origin methylation directions were classified as mixed-sign. Finally, PatMat iDMRs with no overlapping iDMR identified in our analysis were classified as missing.

### Generation of the Pooch Input Data

Pooch uses two intermediate datasets: MethylDataset and DiffDatasetFromMethylDataset. MethylDataset: The lifted methylation levels on T2T-CHM13v2.0 from all trio assembly haplotypes (n = 246) were merged by CpG site position (0-based). The resulting matrix was transposed so that each column represents a genomic position and each row represents a sample haplotype. The first columns contain the sample name and label (Maternal/Paternal or Hap1/Hap2), followed by columns corresponding to CpG positions for the chromosome. If a sample has no value for a given locus, it is filled with NA. This dataset can be generated using the ‘createMethylDataset.py’ script from the Pooch tools. We then manually filtered the resulting dataset to retain only CpG sites with missing values in no more than 5% of sample haplotypes.

DiffDatasetFromMethylDataset: Using the MethylDataset, the difference between the two haplotypes is calculated with the createDiffDatasetFromMethylDataset.py script. The inputs are the MethylDataset and the specified left and right labels, which determine the order of subtraction. For example, if Maternal is set as the left label and Paternal as the right label, the difference is computed as Maternal methylation minus Paternal methylation at each CpG site in each sample and locus. If one of the haplotypes has an NA value at the CpG site, the corresponding locus for that sample is set to 0.

### Pooch model training and test for autosomes

We developed a chromosome-specific binary classifier using elastic net–regularized logistic regression to predict paternal versus maternal haplotype labels from differential methylation features. For each chromosome, samples were split into predefined training and test sets, comprising 80 training samples and 43 test samples. To enforce orientation invariance of the differential features, sign-flipped counterparts were included. Specifically, selected samples were transformed by multiplying all feature columns by (-1), and analogous augmentation was applied to the training set by duplicating each training sample with inverted feature signs and swapped class labels (paternal/maternal). This produced an augmented training matrix for model fitting.

Modeling used a scikit-learn (v1.5.2) (Scikit-learn: Machine Learning in Python, Pedregosa et al., JMLR 12, pp. 2825-2830, 2011) pipeline with select step with SelectFdr(score_func=f_classif, alpha=0.05) and then LogisticRegression (penalty=“elasticnet”, solver=“saga”, max_iter=100000, random_state=42, fit_intercept=False). Note the f_classif computes a one-way ANOVA F statistic. Hyperparameters were optimized by grid search over inverse regularization strength

“model C” : [0.01, 0.1, 1, 10, 100, 1000],

and elastic net mixing parameter

“model l1_ratio“: [0, 0.25, 0.5, 0.75, 1]

So the equation would be :

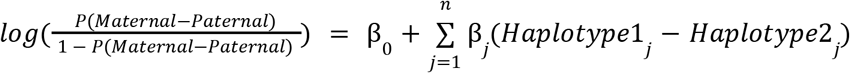

Here, Maternal–Paternal denotes the haplotype configuration, β_0_ is the intercept (set to 0 owing to the symmetric training dataset), and *Haplotype*1*_j_* and *Haplotype*2*_j_* denote the methylation fractions at CpG_j_ on the maternal and paternal haplotypes, respectively.

GridSearchCV was used to identify the optimal hyperparameters using five-fold stratified cross-validation, with negative Brier score and accuracy as evaluation metrics. Note that feature selection of differentially methylated CpGs is entirely restricted to the training set of each fold to avoid data leakage. The final model was refitted using the parameter set that achieved the best predictive performance, as measured by the lowest Brier score. We used GroupKFold(n_splits=5) when splitting the training data into cross-validation folds, ensuring that samples derived from the same original sample were assigned to the same fold.

To obtain a less biased estimate of model performance, we performed nested cross-validation with a five-fold stratified outer loop using the same hyperparameter-tuning procedure.

Out-of-fold predicted probabilities from the outer loop were used to calculate the out-of-fold AUC. Final evaluation on the held-out test set used predicted class probabilities and a decision threshold of 0.5.

### XIST-adjusted elastic net modeling for chromosome X

For chromosome X parent-of-origin prediction, we constructed a balanced training dataset comprising 46 samples, including 23 samples with maternal Xi and 23 samples with paternal Xi. The remaining 13 samples were reserved as an independent test set. To account for XCI-associated methylation variation, we developed an *XIST*-adjusted model. For each sample, differential methylation were calculated at individual CpG sites, together with the differential methylation of the *XIST* promoter (Δ*M_XIST_*). The direction of XIST differential methylation was matched to that of the CpG methylation differences. CpG sites overlapping the *XIST* promoter were excluded from model training. Like for the autosomes, we augmented the dataset by reversing the signs of both CpG methylation differences and *XIST* differential methylation while exchanging the maternal and paternal haplotype labels. The binary response was defined as whether the right haplotype was paternal (*y_i_* = 1) or maternal (*y_i_* = 0), and the corresponding parental-direction variable was defined as ℎ*_i_* = 2*y_i_* − 1.

To distinguish parent-of-origin-associated methylation differences from *XIST*-associated variation, we fitted a CpG-specific linear regression model:

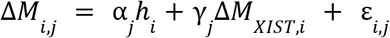

where Δ*Mi_,j_* represents the observed methylation difference at CpG *j* in sample *i*, ℎ*_i_* represents parental direction (−1 or +1), Δ*M_XIST_*_,*i*_ represents *XIST* promoter differential methylation, and α*_j_* and γ*_j_* represent the estimated parent-of-origin- and *XIST*-associated effects, respectively. The regression was fitted by ordinary least squares without an intercept. The *XIST*-associated component was subsequently subtracted from the observed CpG methylation difference:

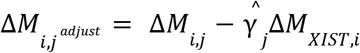

This adjustment was designed to reduce methylation variation linearly associated with *XIST* promoter methylation while retaining the estimated parent-of-origin-associated component. Following *XIST* adjustment, CpG features were screened using ‘SelectFdr(f_classif, alpha=0.05)’ as a feature-selection heuristic. This screening was performed on the adjusted methylation differences independently within each training fold. Importantly, the regression coefficients were estimated exclusively from the training samples within each cross-validation fold and subsequently applied to the corresponding validation samples.

The selected CpG features were subsequently used as input for logistic elastic net classification.

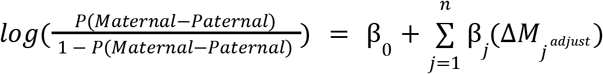

Model coefficients were estimated by minimizing the logistic loss with combined L1 and L2 regularization, without an intercept or an additional independently fitted XIST covariate. Elastic net model fitting, hyperparameter optimization, and nested cross-validation were performed using the same procedures as described for the autosomal models.

### Hapmer and assembly visualization for samples with poor prediction performance

The parent-specific “hapmer” track was generated using Merqury (v1.3), with maternal and paternal k-mers generated by Meryl^65^ (v1.4.1) from the parents’ Illumina data and the personalized diploid assembly, followed by liftover of the BigWig file to the T2T-CHM13v2.0 coordinates using the sample’s chain file.

To demonstrate the incompleteness of chr19 in HG02602, we loaded the PAF file generated during chain file creation and visualized the alignments using SVbyEye^66^ (v0.99.0).

### Model performance using CpGs inside and outside iDMRs

For the “Pooch (iDMR CpGs only)” model, methylation BED files for each sample and haplotype in T2T-CHM13v2.0 coordinates were intersected with PatMat iDMRs using BEDTools. The resulting BED files were used to generate Pooch input datasets with createMethylDataset.py and createDiffDatasetFromMethylDataset.py. To evaluate model performance using CpGs outside PatMat iDMRs, we generated four additional CpG sets by excluding CpGs within PatMat iDMRs and their flanking regions extending 1 bp, 2 kb, 100 kb, or 1 Mb on either side. These exclusions were applied using a custom Python script (script_name.py) to the input dataset (DiffDatasetFromMethylDataset) used to train the autosomal Pooch models after filtering for missing values. Elastic net models were then trained and evaluated as described above, using the same training and held-out test samples across all analyses.

### Haplotype assignment using PatMat

We used the PatMat iDMRs and followed the strategy described in the same study^19^, as outlined below. For each sample and chromosome, we evaluated CpGs located within each annotated iDMR and compared methylation levels between the two haplotypes. An iDMR was considered informative if at least 11 CpGs had non-missing methylation measurements on each haplotype. We then counted CpGs showing a methylation difference of >0.35 between haplotypes and required the absolute difference in the numbers of haplotype-specific CpGs to exceed 10% of the number of informative CpGs on each haplotype. For each retained iDMR, we calculated a weighted methylation score for each haplotype as the mean methylation level across the region multiplied by the fraction of informative CpGs showing haplotype-specific methylation. We summed these scores across all retained maternal- and paternal-biased iDMRs separately for each haplotype. The haplotype with the higher cumulative maternal score was assigned as the maternal haplotype, and the other was assigned as paternal. If the cumulative maternal scores were equal, the haplotype assignment was considered ambiguous. This procedure was performed independently for each chromosome and sample.

### Applying Pooch on independent datasets

For prediction, we developed a dedicated script, prediction.py, which applies the trained chromosome-specific model to the corresponding DiffDatasetFromMethylDataset and calculates prediction probabilities using scikit-learn’s model.predict_proba(X). The input dataset must contain the same CpG features in the same order as those used by the trained model. A compatible dataset can be generated using createMethylDataset.py with the corresponding model. For chromosome X, predict_XIST_adjust.py additionally accepts XIST promoter differential methylation, calculated as described above. The chromosome X model is implemented as a pipeline that uses this value to adjust methylation differences at each CpG site for XCI before prediction. All prediction steps are integrated into the Pooch Snakemake workflow and run automatically for each chromosome (see Implementation of the Pooch Snakemake workflow in Method).

T2T pedigree samples: Four diploid assemblies and the corresponding PacBio HiFi and Oxford Nanopore Technologies (ONT) sequencing datasets were obtained from the pedigree cohort^51^. All sequencing data were converted from BAM to FASTQ format using samtools fastq with the

-T ML,MM option to retain methylation tags. The assemblies and corresponding sequencing data were subsequently analyzed using the Pooch Snakemake pipeline for parent-of-origin haplotype prediction.

KOLF2.1J samples: The KOLF2.1J assembly^52^ was downloaded from https://doi.org/10.5281/zenodo.18777354, along with methylation levels for each CpG site from five neuronal cell lines differentiated from iPSCs, including NGN2-derived neurons, microglia, cortical neurons, astrocytes, and oligodendrocytes, as well as undifferentiated iPSCs. Chain files between each KOLF2.1J haplotype and T2T-CHM13v2.0 were downloaded from https://doi.org/10.5281/zenodo.18777354, and the methylation calling data were lifted over to T2T-CHM13v2.0 coordinates using these chain files. The assemblies and corresponding methylation BED files were subsequently analyzed using the Pooch Snakemake pipeline for parent-of-origin haplotype prediction.

ROB samples: Three diploid assemblies of ROB carriers and their corresponding and ONT sequencing datasets were obtained from a previous study^36^. The assemblies and corresponding sequencing data were subsequently analyzed using the Pooch Snakemake pipeline for parent-of-origin haplotype prediction.

### Gene annotation of iDMRs

We used the T2T-CHM13v2.0 gene annotation (chm13v2.0_RefSeq_Liftoff_v5.3.gtf), downloaded from the CHM13 GitHub repository (https://github.com/marbl/CHM13). A promoter BED file was generated from the gene annotation by defining promoter regions as 250 bp upstream and 100 bp downstream of the TSS of each transcript. The promoter BED file was then intersected with the iDMR BED file to identify iDMRs overlapping promoter regions and to annotate the corresponding genes and transcripts.

### Identification of model coefficient clusters

To identify genomic regions enriched for strong model coefficients, elastic net coefficients were normalized separately for each model by dividing each coefficient by the maximum absolute coefficient, such that the largest absolute coefficient was scaled to one. CpGs with normalized coefficient values ≥0.2 or ≤-0.2 were selected separately for maternal and paternal coefficients.

Within each chromosome, selected CpGs were sorted by genomic position within each chromosome and grouped into clusters when the maximum distance between consecutive CpGs was ≤500 bp. Clusters containing at least three CpGs were retained as strong coefficient clusters.

### Region-based haplotype-by-XIST interaction analysis

To assess the effects of parental haplotype and XIST promoter methylation separately at putative ICRs on chromosome X, we obtained previously reported putative ICRs on chromosome X from [humanicr.org](https://humanicr.org/)^45^. Their coordinates were lifted over to T2T-CHM13v2.0 using CrossMap^64^ (v0.7.3) and the GRCh38-to-CHM13v2.0 chain file from the [CHM13 GitHub repository](https://github.com/marbl/CHM13). Mean methylation levels within each region were then calculated separately for each haplotype in a set of 50 samples balanced for XCI status, as described above.

Then, we performed region-based interaction analyses between parental haplotype and *XIST* promoter differential methylation. We fitted a linear mixed-effects model to evaluate the effects of parental haplotype, *XIST* promoter differential methylation, and their interaction on regional methylation levels, while accounting for repeated measurements from the same sample using a random intercept:

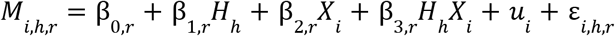

where *M_i_*_,ℎ,*r*_ represents the coverage-weighted methylation proportion for haplotype ℎ of sample *i* at region *r*, *H*_ℎ_ represents parental haplotype (maternal or paternal), *X_i_* represents XIST promoter differential methylation for sample *i*, and β_1,*r*_, β_2,*r*_, and β_3,*r*_ represent the haplotype, *XIST* differential methylation, and interaction effects, respectively. The sample-specific random intercept (*u_1_*) accounts for within-sample correlation between maternal and paternal measurements.

Linear mixed-effects models were fitted using ‘MixedLM’ from the Python ‘statsmodels’ package (v0.15.0) (’statsmodels.formula.api.mixedlm’) (Seabold, Skipper, and Josef Perktold. “statsmodels: Econometric and statistical modeling with python.” Proceedings of the 9th Python in Science Conference. 2010.). P-values for each fixed-effect term were adjusted for multiple testing using the Benjamini–Hochberg FDR procedure, implemented with ‘statsmodels.stats.multitest.multipletests(method=“fdr_bh”)’. FDR correction was performed separately for each fixed-effect term (haplotype, *XIST* differential methylation, and their interaction) across all tested regions.

### Chromosome-wide CpG-level interaction analysis

To investigate parent-of-origin-associated methylation and its interaction with XCI at individual CpG sites across chromosome X, we performed chromosome-wide linear regression analysis using paired maternal and paternal methylation data from 50 samples balanced for XCI status.

For each sample, differential methylation at each CpG site was calculated as maternal minus paternal methylation (Mat − Pat, Δ*M_s_*). We then fitted a separate linear regression model for each CpG site, using CpG differential methylation (Δ*M_s_*) as the response variable and XIST promoter differential methylation (Δ*XIST_s_*) as the predictor.

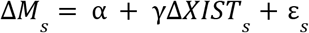

The model intercept (α) represented the estimated parental haplotype methylation difference when XIST promoter differential methylation was zero, whereas the slope (γ) represented the association between XIST promoter differential methylation and the parental methylation difference at each CpG site. Only CpG sites with paired methylation measurements in all 50 samples and sufficient variation in XIST promoter differential methylation were included.

Regression coefficients and standard errors were calculated using NumPy (v2.4.6), and two-sided P-values for the intercept and slope were calculated using the Student’s t distribution in SciPy (v1.17.1). P-values were adjusted separately for each term across all tested CpGs using the Benjamini–Hochberg FDR correction implemented in statsmodels (v0.15.0).

### Comparison of PAR1 methylation patterns in XY samples

Methylation calls from the paternal Y chromosome were first lifted over from personalized genome coordinates to the T2T-CHM13v2.0 chromosome Y reference using CrossMap^64^ (v0.7.0) and the corresponding chain files. The coordinates were then lifted over to T2T-CHM13v2.0 chromosome X using CrossMap and a custom chain file generated from the CHM13 chromosome X and Y sequences, as described above, to generate pseudo-paternal X (pseudo-pX) methylation profiles for 64 XY HPRC samples. To calculate mean methylation levels at the DHRSX iDMR (chrX:1,937,651–1,938,190), we extracted overlapping CpG methylation calls using BEDTools intersect (v2.31.1) and summed methylated read counts and total read coverage using BEDTools merge. The methylation fraction for each sample and haplotype was calculated by dividing the total methylated read count by the total read coverage across the region. Differences in methylation fractions between the maternal X (mX) and pseudo-pX haplotypes were evaluated using paired t-tests implemented with scipy.stats.ttest_rel(). The analysis included 63 of the 64 XY samples. HG02514 was excluded because no maternal X chromosome CpG methylation calls were available within the analyzed region

### Implementation of the Pooch Snakemake workflow

The entire Pooch pipeline was built using Snakemake^67^ (v7.32.4). Within the Pooch pipeline, the input FASTQ files and personalized diploid assemblies are automatically aligned, followed by methylation calling, chain file generation, liftover to T2T-CHM13v2.0, parent-of-origin prediction, and reporting of assembly errors when present. The detailed pipeline is described in Fig. 5 and below.

Methylation tag check: Pooch accepts FASTQ files, BAM files aligned to a personalized genome, or methylation calls in personalized genome coordinates. FASTQ and BAM inputs are checked for MM and ML tags, and only datasets containing these methylation tags proceed to subsequent analyses.

Preprocessing assemblies of haplotype: Pooch accepts either diploid assembly (.fasta) or pairs of haploid assemblies as personalized genome input. When haploid assemblies are provided, a diploid assembly is generated by concatenating the two haploid assemblies prior to mapping.

Align: To improve alignment accuracy in repetitive regions, Pooch uses Winnowmap2 as default mapper. The repetitive k-mers were identified from the personalized diploid assembly using Meryl. Specifically, 15-mer frequencies were counted using ‘meryl count’ to generate a k-mer database, and repetitive k-mers were extracted by retaining k-mers with a distinct frequency threshold greater than 0.9998. Reads were aligned to the reference genome using Winnowmap2 with the platform-specific preset (map-ont or map-pb), retaining methylation-related tags were applicable (-y -Y). For the minimap2 option, miniamp2 was run with –MD, –cs, -Y and -y, and the platform-specific prest (-ax map-ont or map-bp). Then only primary alignments remained after using ‘samtools view -F 0×104’.

Chaining: As described above.

Methylation calling: Methylation calling is done by different tools depending on the input file platform, modkit for ONT, pb-cpg-tools for HiFi.

Liftover: As described above.

Chromosome assignment: Using the chain files, we extracted the mapping regions between T2T-CHM13v2.0 and each haplotype assembly. Mapping regions were then aggregated by contig and chromosome, and only contig–chromosome pairs with a minimum total alignment length of 10 kb were retained. For each chromosome, the contig with the longest total alignment length was selected and reported.

Sex check: Prediction for chromosome X was performed only for samples with two X chromosomes. Based on the chromosome assignment results, samples were classified as male if chromosome Y was present in the chromosome assignment file and as female otherwise.

Classification: For each chromosome, a MethylDataset was generated from the corresponding methylation BED files, retaining only the CpGs used by the trained chromosome-specific model.

A DiffDataset was then generated by calculating methylation differences according to the user-defined haplotype configuration. For example, “haplotype1” and “haplotype2” could be designated as hap1 and hap2, respectively, with differences calculated as hap1 − hap2. The trained models were applied using prediction.sh for autosomes and predict_XIST_adjust.py for chromosome X to generate the final parent-of-origin haplotype assignment file.

### Subsampling analysis

We applied the same pipeline described above while subsampling the training set to 10, 15, 20, 25, 30, 40, 50, 60, 70, or 80 samples to determine the minimum training-set size required to achieve high prediction accuracy. Only the training samples were subsampled, whereas the test set was kept unchanged across all analyses. This design allowed test accuracy to be directly compared with the results shown in Fig. 2.

### Computational resource requirements

To assess the computational resources required for the prediction step, we used the methylation-calling BED files for the maternal and paternal haplotypes of HG002 for autosomes and HG01081 for chrX, lifted over to T2T-CHM13v2.0 coordinates. We generated the input datasets for Pooch using createMethylDataset.py and createDiffDatasetFromMethylDataset.py, and then performed parent-of-origin prediction using predict.py and predict_XIST_adjust.py. Computational resource usage was measured using GNU time (time -v). Wall-clock time was recorded as the total elapsed execution time, CPU time was calculated as the sum of user and system CPU times, and peak memory usage was estimated from the maximum resident set size (RSS).

## Data availability

The comprehensive iDMRs reported in this paper along with their corresponding genome browser tracks are available at https://github.com/marbl/Pooch. The 123 diploid assemblies generated with Verkko v2.2.1 using both trio and Hi-C data for phasing are available in the HPRC S3 bucket (https://s3-us-west-2.amazonaws.com/human-pangenomics/index.html?prefix=publications/Poo ch_2026/). This archive contains the original assembly files used in this study.

## Code availability

All Pooch code is available from https://github.com/marbl/Pooch, along with links to tutorials, pre-trained models, and all scripts used in this study.

## Acknowledgements

We would like to acknowledge the National Human Genome Research Institute (NHGRI) for funding the following grants supporting the creation of the human pangenome reference: UM1HG010971, U41HG010972, U01HG013760, U01HG013755, U01HG013748, U01HG013744, R01HG011274, and the Human Pangenome Reference Consortium (BioProject ID: PRJNA730823). J.K. was supported by Aligning Science Across Parkinson’s MJFF-029775 through the Michael J. Fox Foundation for Parkinson’s Research (MJFF). This research was supported in part by the Intramural Research Program of the National Institutes of Health (NIH). The contributions of the NIH author(s) are considered Works of the United States Government. The findings and conclusions presented in this paper are those of the author(s) and do not necessarily reflect the views of the NIH or the U.S. Department of Health and Human Services. This work utilized the computational resources of the NIH HPC Biowulf cluster (https://hpc.nih.gov).

## Author Information

### Contributions

A.M.P. conceived of and supervised the work. J.K. performed model fitting and validation, wrote the main execution script, developed the pipelines, and drafted the paper. B.P. contributed to the conceptualization of the study and developed the original Python modules. J.B. and B.D.P. performed exploratory pilot analyses. N.F.H. generated HPRC assembly alignments and chain files. M.A.W. contributed to the interpretation of chrX methylation patterns. S.K. and A.R. contributed to analyses and project direction. All authors reviewed, edited, and approved the final version of the manuscript.

## Ethics declarations

### Consortia

Human Pangenome Reference Consortium (affiliations in Supplemental list of authors)

Derek Albracht, Ivan A. Alexandrov, Jamie Allen, Alawi A. Alsheikh-Ali, Nicolas Altemose, Casey Andrews, Dmitry Antipov, Lucinda Antonacci-Fulton, Alexander Arguello, Mobin Asri, Marcelo Ayllon, Jennifer R. Balacco, Floris P. Barthel, Edward A. Belter Jr, Halle D. Bender, Andrew P. Blair, Davide Bolognini, Katherine E. Bonini, Christina Boucher, Guillaume Bourque, Silvia Buonaiuto, Shuo Cao, Andrew Carroll, Ann M. Mc Cartney, Monika Cechova, Mark J.P. Chaisson, Pi-Chuan Chang, Xian Chang, Jitender Cheema, Haoyu Cheng, Claudio Ciofi, Hiram Clawson, Sarah Cody, Vincenza Colonna, Holland C. Conwell, Robert Cook-Deegan, Mark Diekhans, Maria Angela Diroma, Daniel Doerr, Zheng Dong, Danilo Dubocanin, Richard Durbin, Jana Ebler, Evan E. Eichler, Jordan M. Eizenga, Parsa Eskandar, Eddie Ferro, Anna-Sophie Fiston-Lavier, Sarah M. Ford, Willard W. Ford, Giulio Formenti, Adam Frankish, Mallory A. Freeberg, Qichen Fu, Stephanie M. Fullerton, Robert S. Fulton, Shenghan Gao, Yan Gao, Gage H. Garcia, Obed A. Garcia, Joshua M.V. Gardner, Shilpa Garg, Erik Garrison, Nanibaa’ A. Garrison, John E. Garza, Margarita Geleta, Mohammadmersad Ghorbani, Tina A. Graves-Lindsay, Richard E. Green, Carol W. Greider, Cristian Groza, Bida Gu, Andrea Guarracino, Melissa Gymrek, Maximilian Haeussler, Leanne Haggerty, Ira M. Hall, Nancy F. Hansen, Yue Hao, Mohammad Amiruddin Hashmi, David Haussler, Prajna Hebbar, Peter Heringer, Glenn Hickey, Todd L. Hillaker, S. Nakib Hossain, Neng Huang, Sarah E. Hunt, Toby Hunt, Alexander G. Ioannidis, Nafiseh Jafarzadeh, Nivesh Jain, Erich D. Jarvis, Maryam Jehangir, Juan Jiang, Eimear E. Kenny, Juhyun Kim, Bonhwang Koo, Sergey Koren, Milinn Kremitzki, Charles H. Langley, Ben Langmead, Heather A. Lawson, Daofeng Li, Heng Li, Ronghan Li, Wen-Wei Liao, Jiadong Lin, Tianjie Liu, Glennis A. Logsdon, Ryan Lorig-Roach, Jonathan LoTempio Jr, Hailey Loucks, Jane E. Loveland, Jianguo Lu, Shuangjia Lu, Julian K. Lucas, Walfred Ma, Juan F. Macias-Velasco, Kateryna D. Makova, Maximillian G. Marin, Christopher Markovic, Tobias Marschall, Franco L. Marsico, Fergal J. Martin, Mira Mastoras, Capucine Mayoud, Brandy McNulty, Jack A. Medico, Julian M. Menendez, Karen H. Miga, Anna Minkina, Matthew W. Mitchell, Saswat K. Mohanty, Younes Mokrab, Jean Monlong, Shabir Moosa, Avelina Moreno-Ochando, Shinichi Morishita, Jonathan M. Mudge, Katherine M. Munson, Njagi Mwaniki, Nasna Nassir, Chiara Natali, Shloka Negi, Lingbin Ni, Adam M. Novak, Faith Okamoto, Keisuke K. Oshima, Pilar N. Ossorio, Chie Owa, Sadye Paez, Benedict Paten, Clelia Peano, Adam M. Phillippy, Brandon D. Pickett, Laura Pignata, Nadia Pisanti, David Porubsky, Pjotr Prins, Timofey Prodanov, Anandi Radhakrishnan, T. Rhyker Ranallo-Benavidez, Brian J. Raney, Mikko Rautiainen, Alessandro Raveane, Andreas Rechtsteiner, Luyao Ren, Arang Rhie, Fedor Ryabov, Samuel Sacco, Farnaz Salehi, Michael C. Schatz, Laura B. Scheinfeldt, Aarushi Sehgal, William E. Seligmann, Mahsa Shabani, Kishwar Shafin, Shadi Shahatit, Ruhollah Shemirani, Vikram S. Shivakumar, Swati Sinha, Jouni Sirén, Linnéa Smeds, Steven J. Solar, Marco Sollitto, Nicole Soranzo, Andrew B. Stergachis, Marie-Marthe Suner, Yoshihiko Suzuki, Arda Söylev, Ahmad Abou Tayoun, Jack A.S. Tierney, Chad Tomlinson, Francesca Floriana Tricomi, Mohammed Uddin, Matteo Tommaso Ungaro, Rahul Varki, Flavia Villani, Ivo Violich, Mitchell R. Vollger, Brian P. Walenz, Charles Wang, Lisa E. Wang, Ting Wang, Aaron M. Wenger, Conor V. Whelan, Zilan Xin, Zheng Xu, Kai Ye, DongAhn Yoo, Wenjin Zhang, Ying Zhou, Xiaoyu Zhuo, Giulia Zunino

