## Supplementary Fig. for "Parent-of-origin classification for diploid genomes using Pooch"

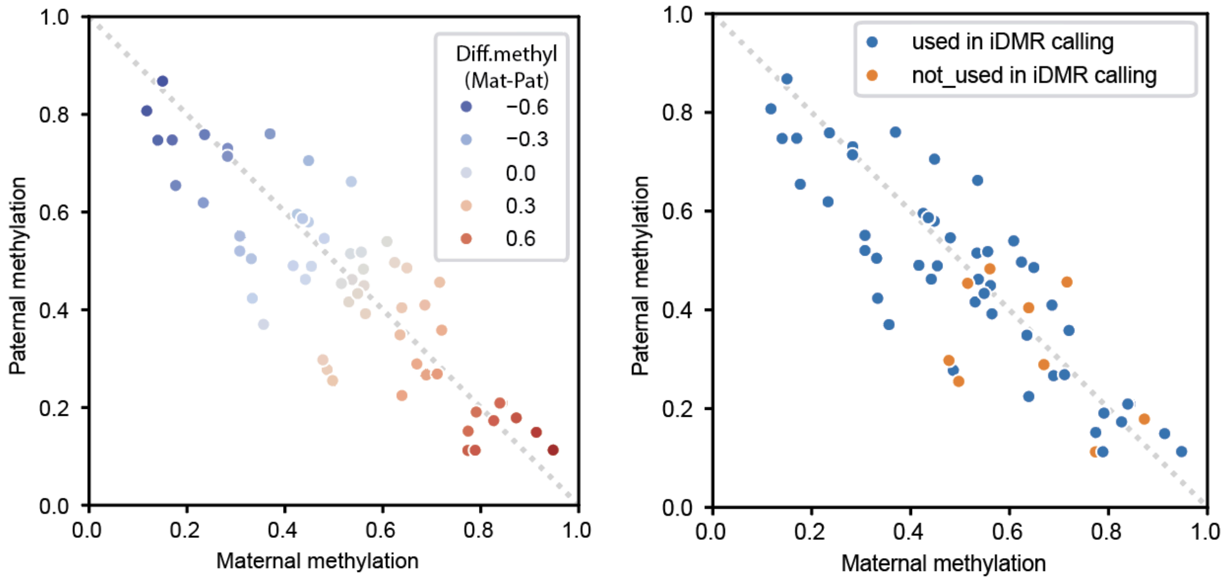


**Supplementary Fig. 1: Mean methylation levels at the XIST promoter by parental haplotype.**

Mean methylation levels at the XIST promoter are plotted for maternal and paternal haplotypes for 59 Female samples. Points are colored by differential methylation between haplotypes (maternal − paternal, left) or by whether the sample was used for iDMR calling (right).

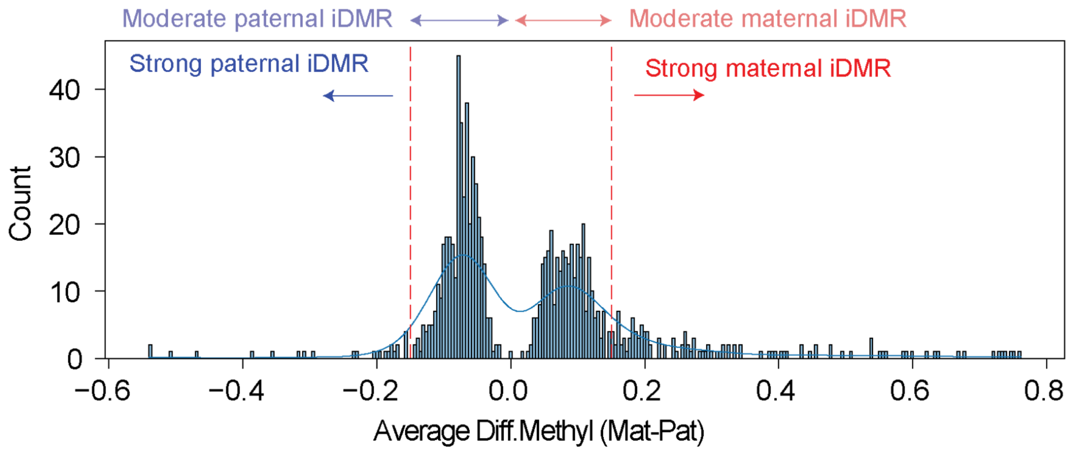


**Supplementary Fig. 2: Distribution of differential methylation levels across iDMRs.**

The x-axis represents the difference in methylation levels between maternal and paternal haplotypes (maternal − paternal). Red dashed lines at −0.15 and 0.15 mark the boundaries between strong and moderate iDMRs. Strong iDMRs have an absolute methylation difference ≥ 0.15, whereas moderate iDMRs have an absolute methylation difference < 0.15.

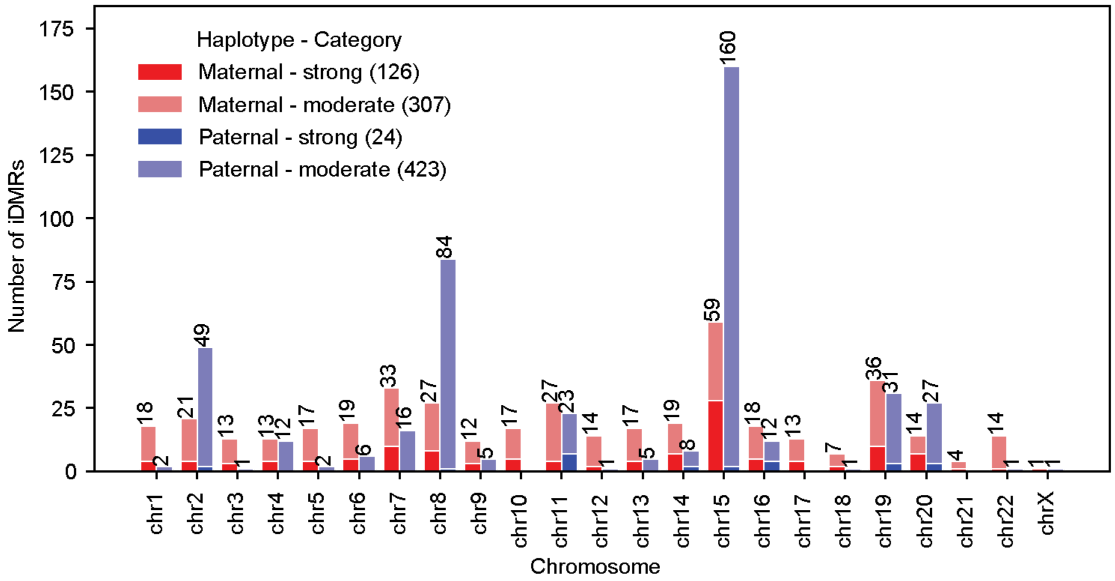


**Supplementary Fig. 3: Number of iDMRs by chromosome, parental hypermethylated haplotype, and methylation category.**

The stacked bar plot shows the number of strong and moderate iDMRs for each chromosome, with maternally and paternally hypermethylated iDMRs displayed side by side.

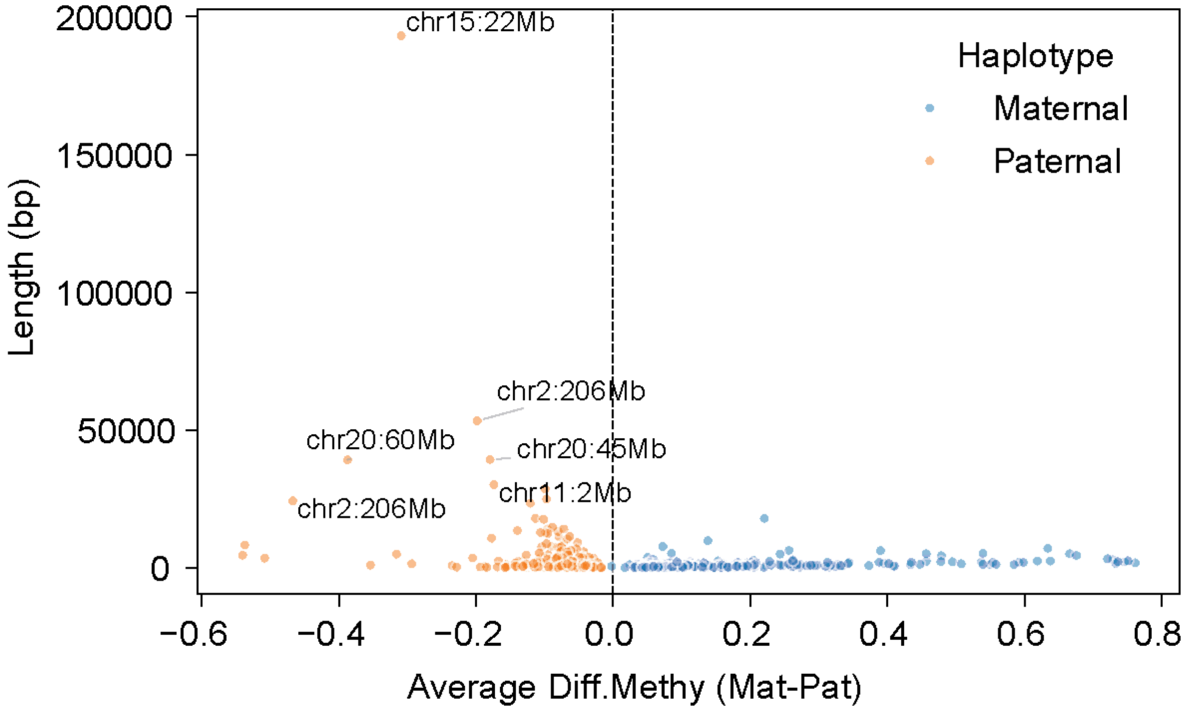


**Supplementary Fig. 4: Relationship between differential methylation levels and iDMR lengths.**

The scatter plot shows the differential methylation between maternal and paternal haplotype (Mat - Pat) on the x-axis and iDMR length on the y-axis. Labels indicate the genomic coordinates of iDMRS based on the T2T-CHM13 reference genome.

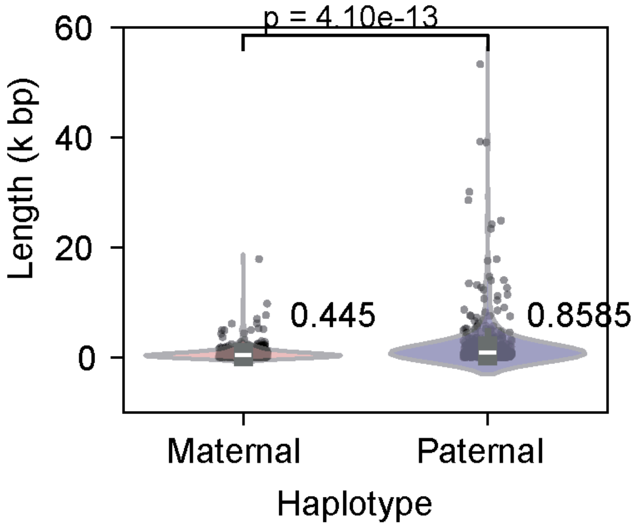


**Supplementary Fig. 5: Length distributions of miDMRs and piDMRs after excluding the longest piDMR.**

Length distributions of maternally and paternally hypermethylated iDMRs. The P-value was calculated using the Wilcoxon rank-sum test after excluding the longest piDMR.

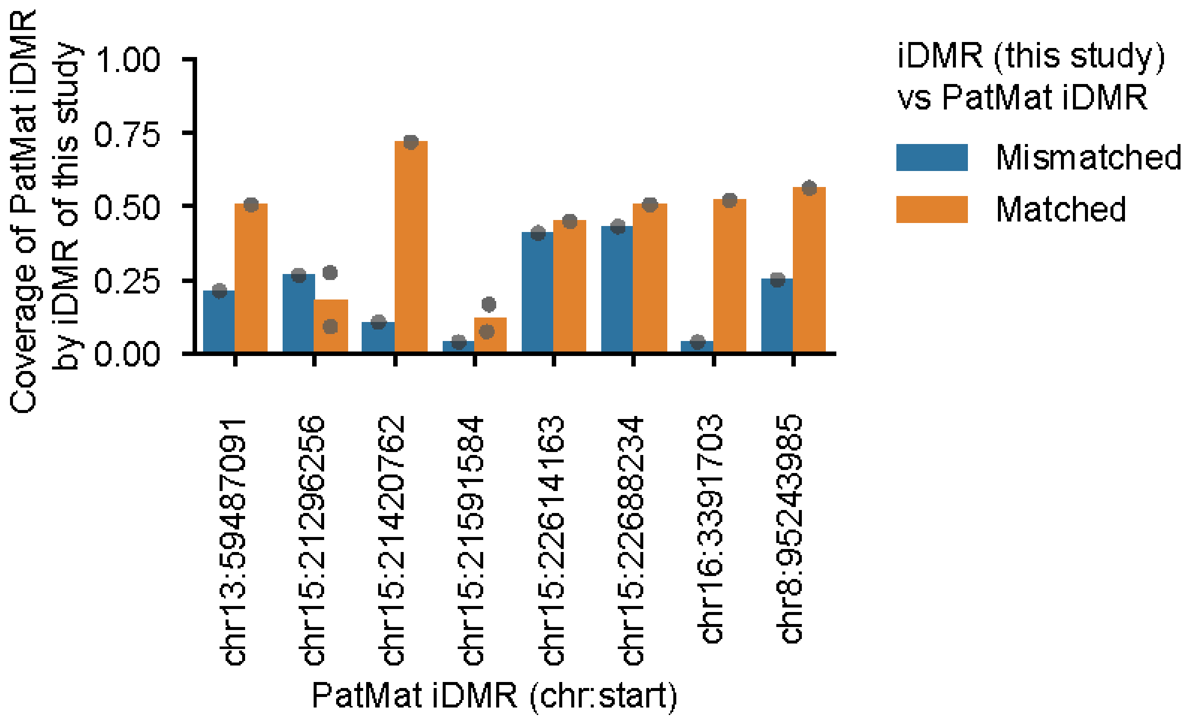


**Supplementary Fig. 6: Coverage of PatMat iDMR of mixed-sign iDMRs.**

Coverage of regions overlapping between PatMat iDMRs and the iDMR list discovered in this study. Consistent directionality shows high coverage (orange), whereas discordant directionality shows low coverage (blue) for each PatMat iDMR.

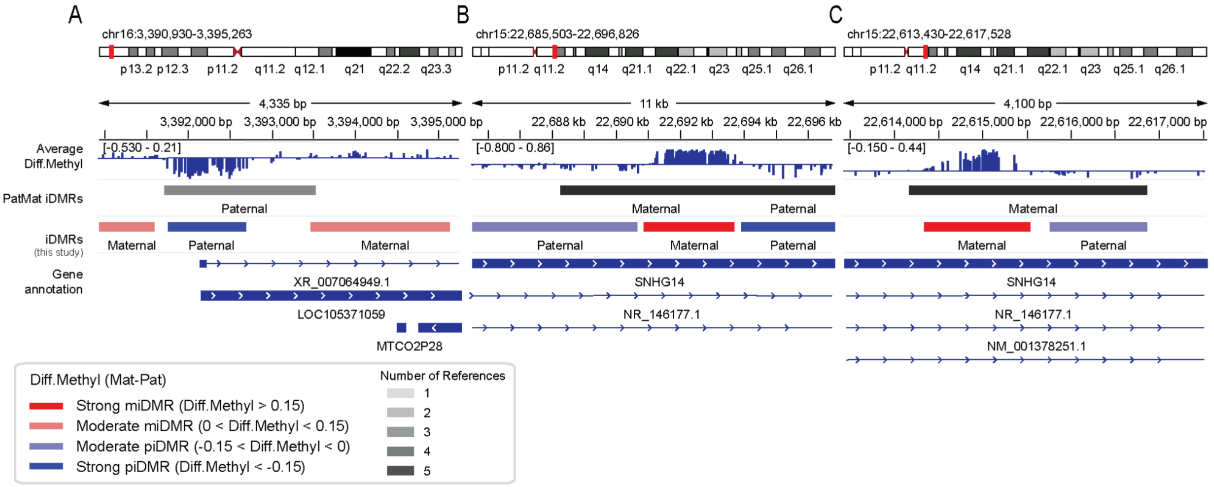


**Supplementary Fig. 7: Genomic views of mixed-sign iDMRs.**

Representative examples of mixed-sign iDMRs are shown. The “Average Diff.Methyl” track displays the mean methylation difference between maternal and paternal haplotype (Mat - Pat). The “PatMat iDMRs” track shows regions identified in previous studies, with colors indicating the number of supporting references. The “iDMRs (This study)” track displays iDMRs identified in this study, colored according to their classification categories. The “Gene Annotation” track shows gene and transcripts based on the latest T2T-CHM13 gene annotation (JHU RefSeq + Liftoff v5.3).

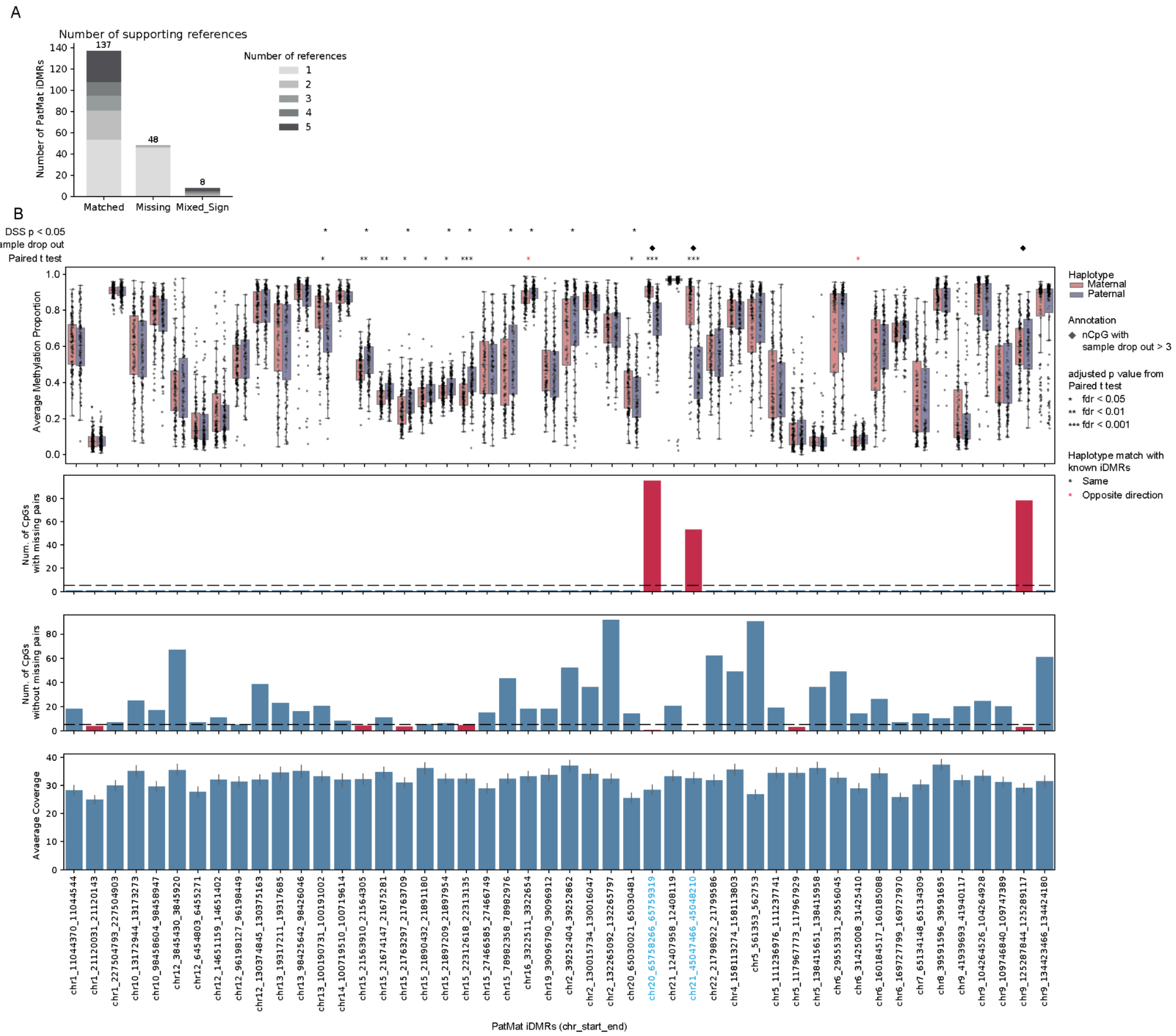


**Supplementary Fig. 8: Analysis of PatMat iDMRs not recovered in this study.**

(A) Number of PatMat iDMRs stratified by the number of independent supporting studies. The x-axis indicates the overlap categories between iDMRs identified in this study and PatMat iDMRs. (B) The panels show, from top to bottom, mean methylation levels of maternal and paternal haplotypes across PatMat iDMRs not recovered in this study, the number of CpG sites with missing paired data in at least one sample, the average number of CpG sites retained per sample after excluding those with missing data, and the mean CpG coverage within each iDMR. Asterisks in the top panel indicate iDMRs detected by DSS using a less stringent P-value threshold of 0.05 instead of 0.001, while diamonds indicate regions with coverage dropout affecting more than three CpG sites on both haplotypes in at least two samples. The third row, labeled “Paired t-test,” indicates significant methylation differences between parental haplotypes based on paired t-tests of region-level mean methylation rather than individual CpG measurements. In the x-axis labels, iDMRs supported by more than one independent study are highlighted in blue.

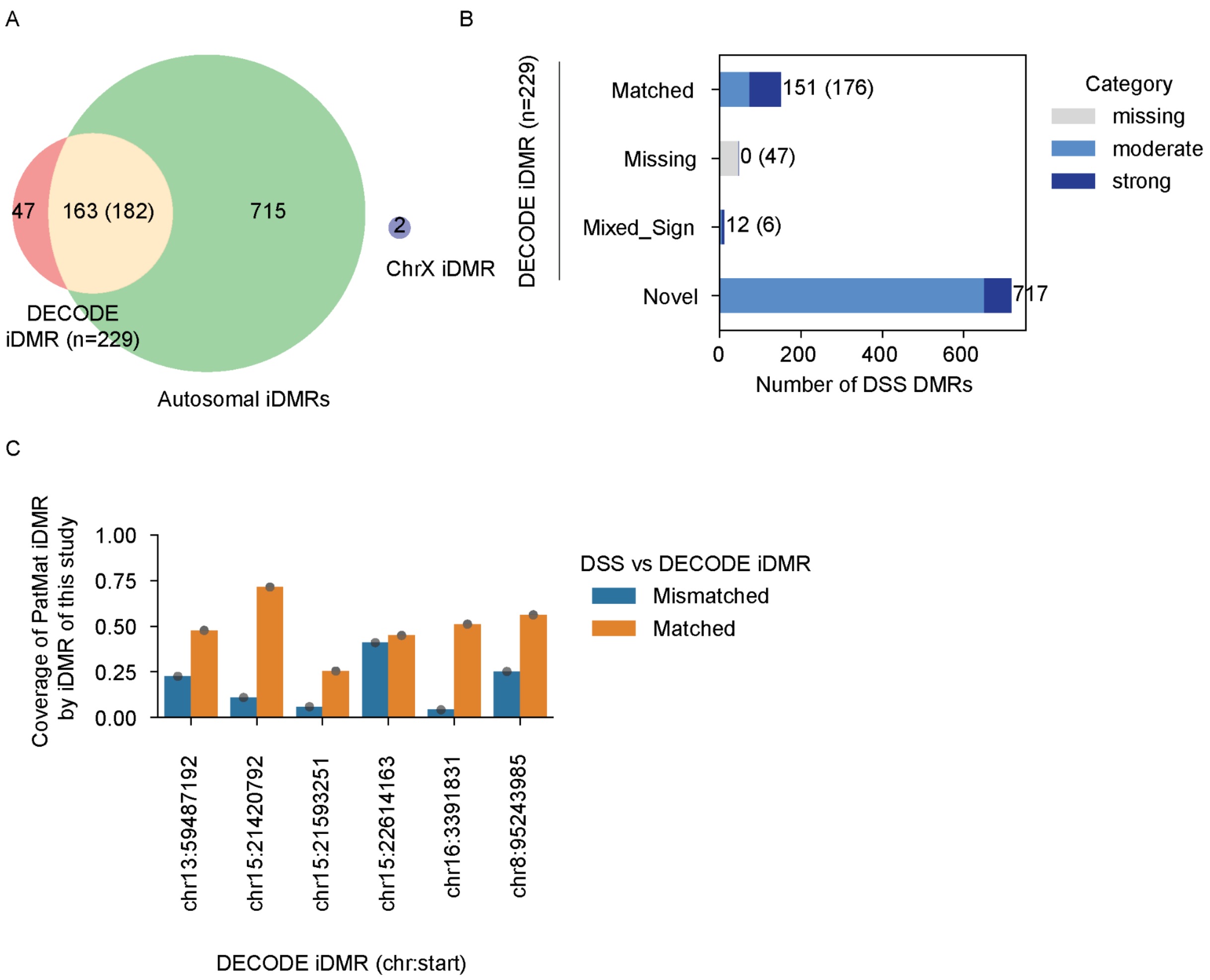


**Supplementary Fig. 9: Comparison of iDMRs identified in this study with the deCODE iDMR catalog.**

(A) Venn diagram showing the overlap between DECODE iDMRs and those identified in this study. iDMRs on chrX are shown separately. Numbers indicate iDMRs identified in this study, with unique DECODE iDMR counts in parentheses. (B) Number of iDMRs identified in this study and those overlapping the 229 DECODE iDMRs. Numbers indicate iDMRs identified in this study, with unique DECODE iDMR counts in parentheses. Colors indicate differential methylation categories (strong or moderate), with gray representing missing iDMRs. The y-axis shows overlap categories: “Matched” indicates overlapping iDMRs with concordant hypermethylated parental haplotypes; “Mixed-sign” indicates DECODE iDMRs overlapping multiple iDMRs identified in this study with opposing methylation directions; and “Missing” indicates DECODE iDMRs with no overlap. (C) Coverage of regions overlapping between PatMat iDMRs and the iDMR list discovered in this study. Consistent directionality shows high coverage (orange), whereas discordant directionality shows low coverage (blue) for each DECODE iDMR.

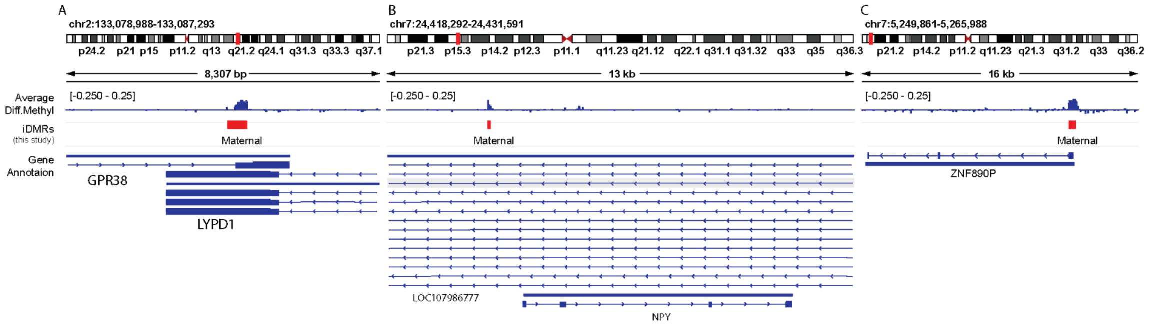


**Supplementary Fig. 10: Genomic views of candidate imprinted loci associated with strong iDMRs.**

(A) Promoter region of *GPR39* and *LYPD1*. (B) Region approximately 1 kb upstream of the *NPY* transcription start site. (C) Promoter region of *ZNF890P*.

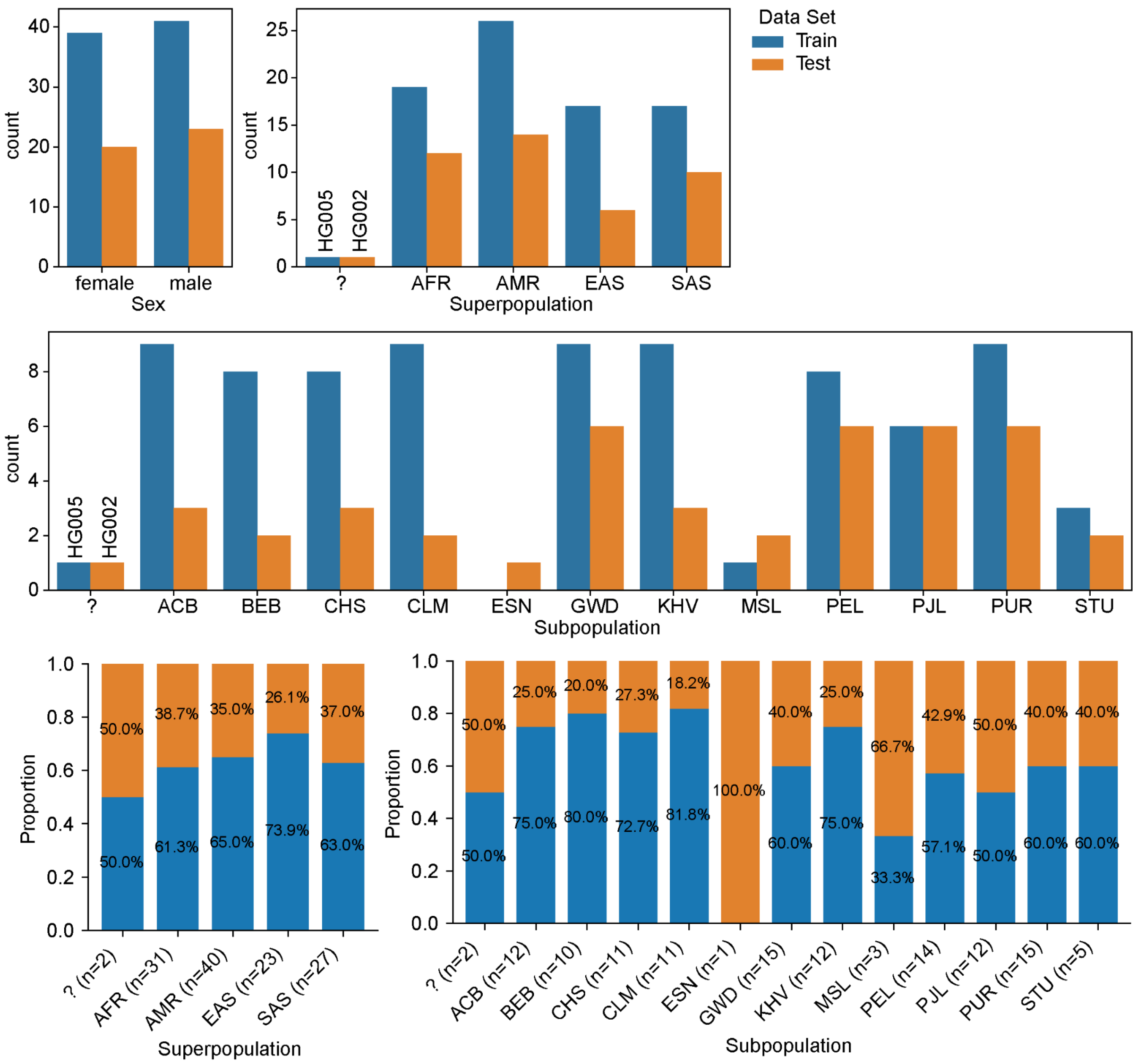


**Supplementary Fig. 11: Sex, superpopulation and subpopulation distribution of training and test samples.**

The numbers of training and test samples are stratified by sex, superpopulation, and subpopulation. Samples with “?” annotations in the HPRC metadata for superpopulation or subpopulation are labeled by sample name in the plot.

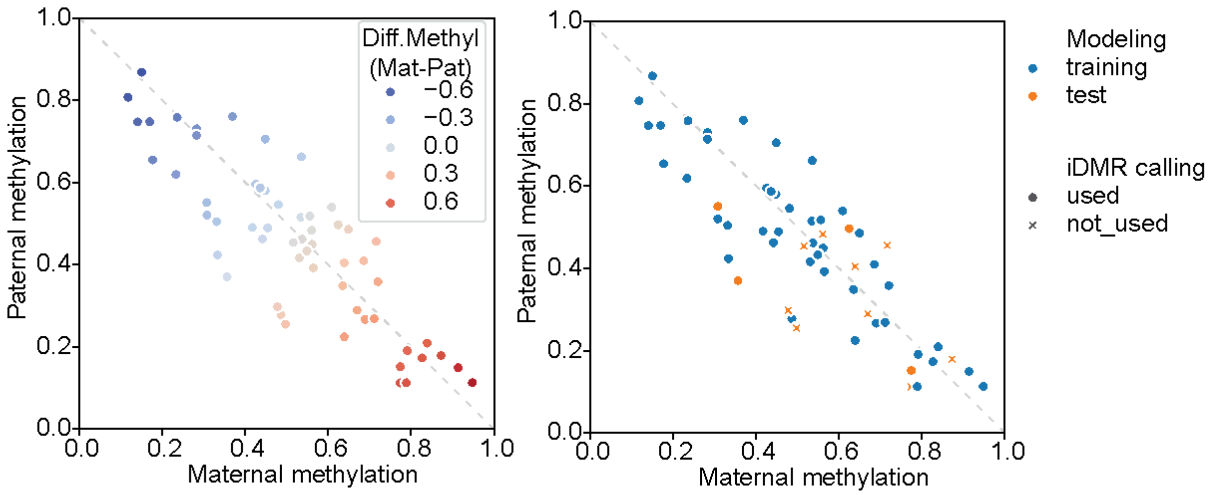


**Supplementary Fig. 12: Mean methylation levels at the XIST promoter for each haplotype.**

For each sample and haplotype, methylated read counts and total read coverage were summed across CpG sites within the XIST promoter (chrX:72,285,969-72,286,319). The mean methylation level was then calculated by dividing the total number of methylated reads by the total read coverage.

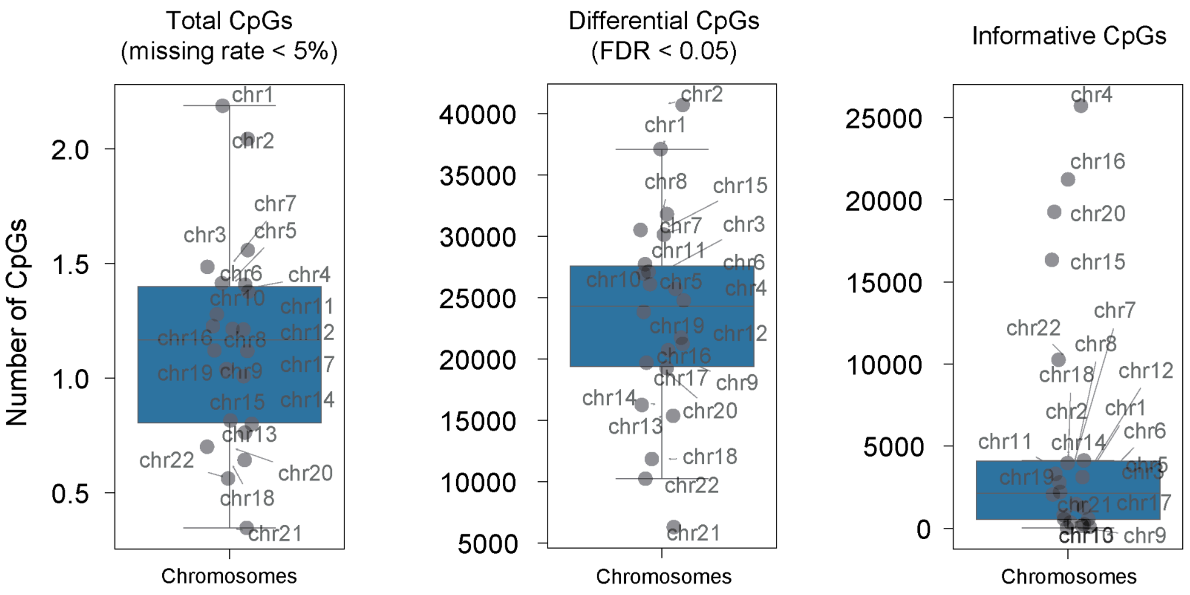


**Supplementary Fig. 13: Changes in the number of selected CpGs during model training**

Total CpGs include all CpGs successfully lifted over to CHM13 with a missing rate <5%. Differential CpGs are CpGs that passed the FDR-adjusted ANOVA test during cross-validation using the parameter set that yielded the best model performance. Predictive CpGs are CpGs with non-zero coefficients in the final models.

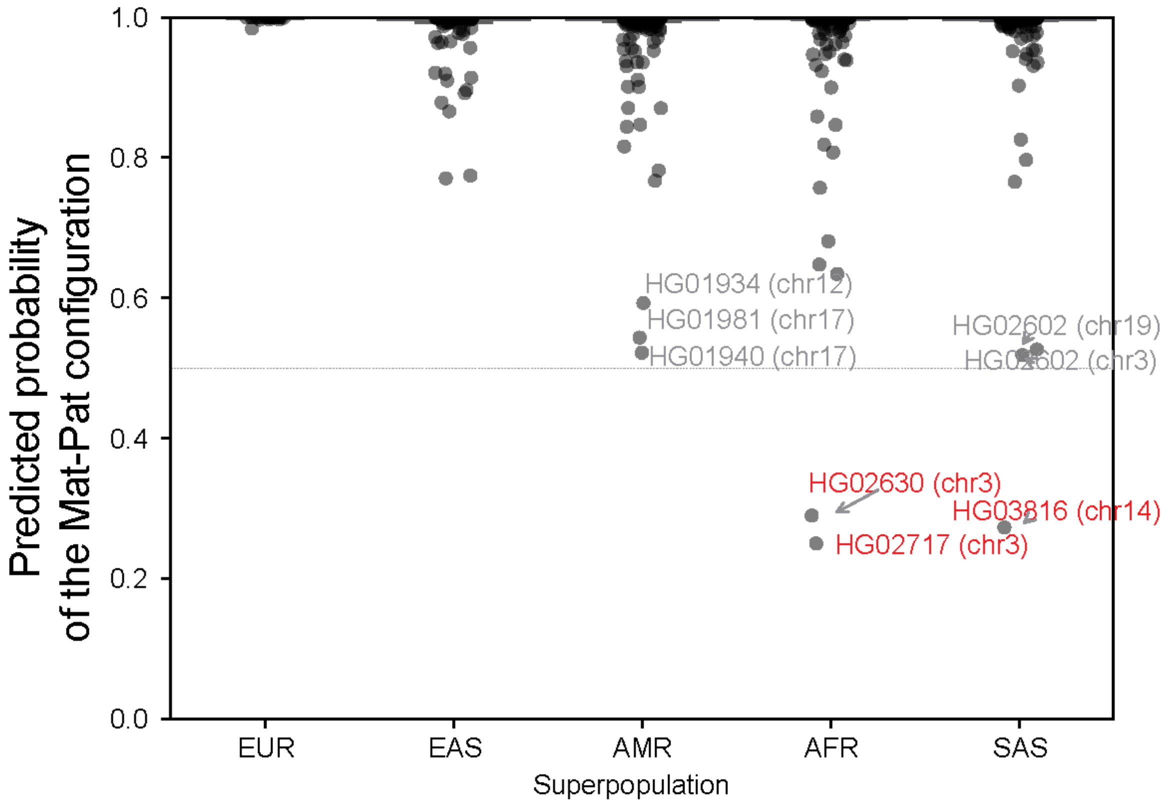


**Supplementary Fig. 14: Pooch prediction stratified by superpopulation and subpopulation**

The results are the same as those shown in Fig. 2B, with samples regrouped by superpopulation. Samples with prediction probabilities below 0.6 are labeled in gray, and those with probabilities below 0.5 are labeled in red. In this figure, HG002 was assigned to the European (EUR) superpopulation.

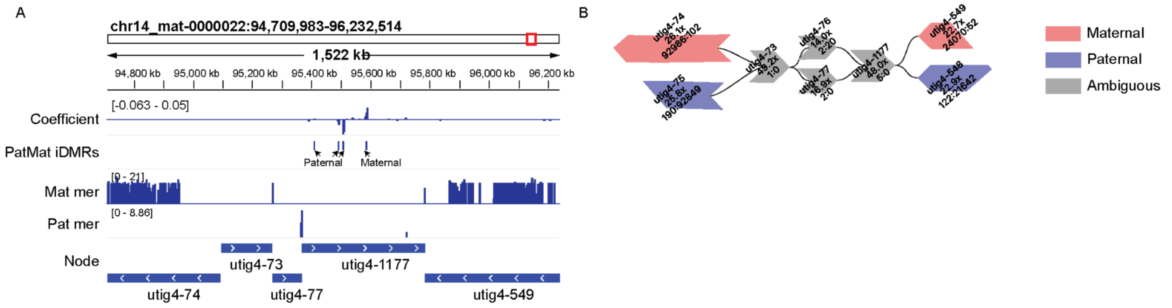


**Supplementary Fig. 15: A homozygous region at an imprinted locus on chromosome 14 of HG03816.**

(A) Pooch model coefficients and parent-specific k-mers on the maternal chromosome 14 haplotype of HG03816, which showed poor haplotype assignment by Pooch. Histograms of Mat- or Pat-specific k-mers indicate an extended region of homozygosity. Assembly node alignments are shown below. (B) Assembly graph topology of HG03816 at the locus shown in (A), highlighting homozygous nodes.

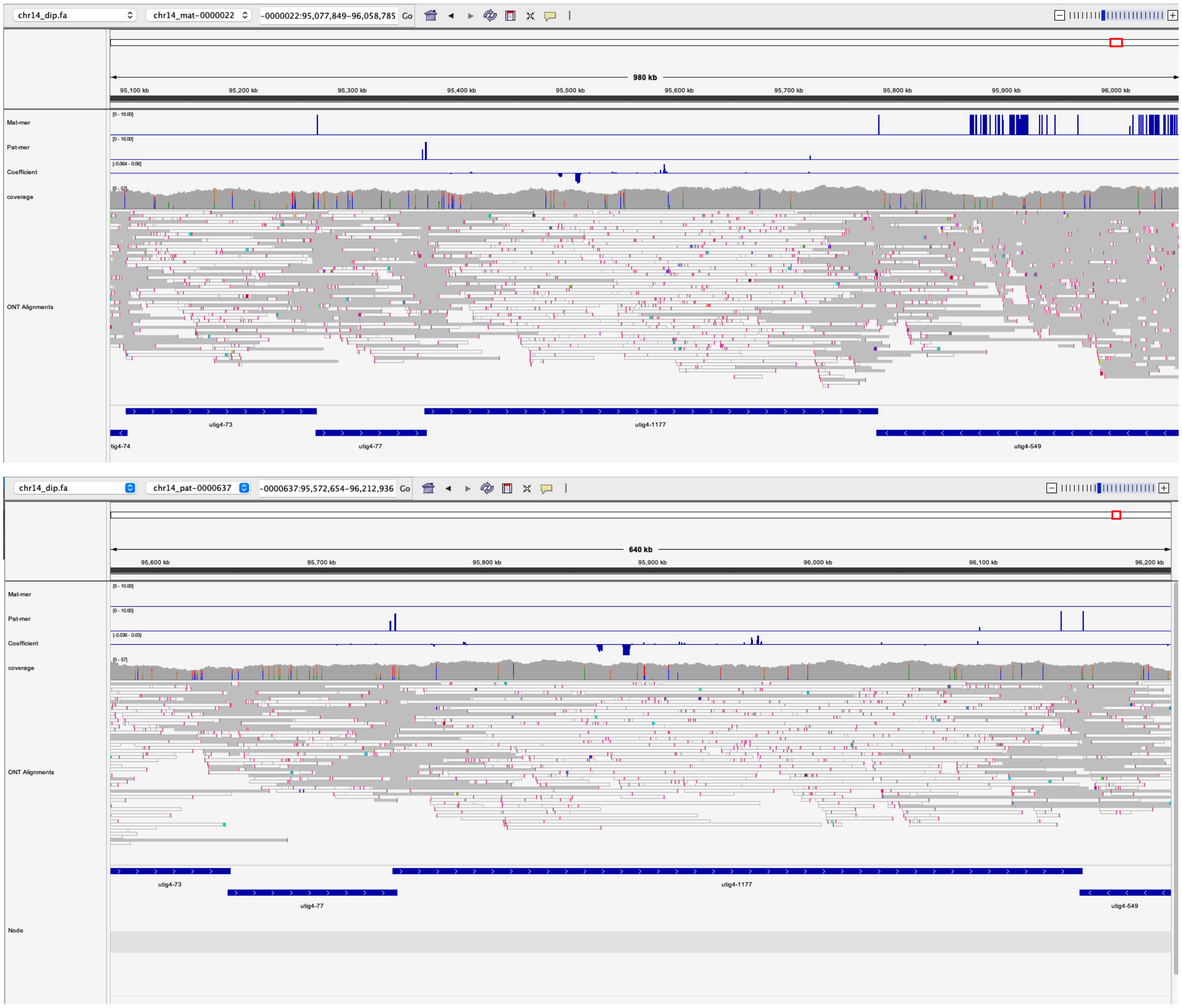


**Supplementary Fig. 16: ONT read alignments of HG03186 at the region shown in Fig. 2C and 2D.**

ONT read alignments to the personalized maternal (top) and paternal (bottom) assemblies, with Mat-mer, Pat-mer, coefficients, coverage, read alignments, and nodes shown. The region around the long, homogeneous node utig4-1177 has low mapping quality (MAPQ = 0) and is displayed in light gray in the IGV plot.

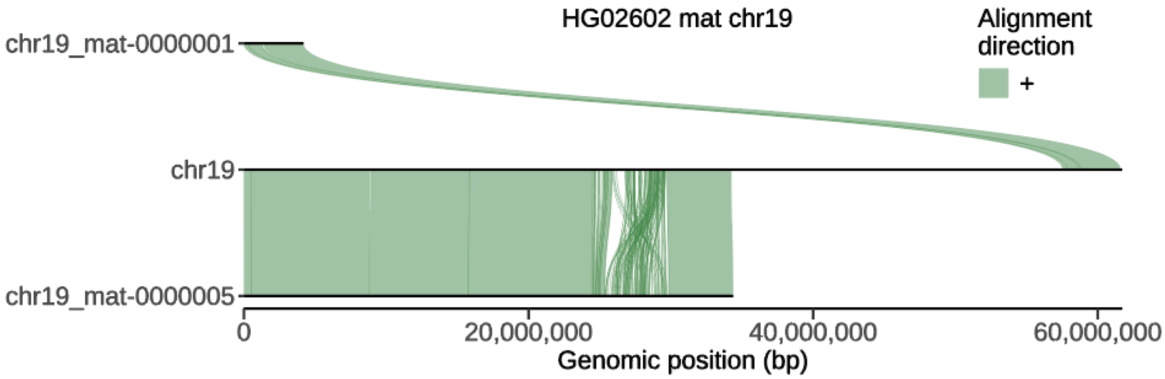


**Supplementary Fig. 17: Synteny between CHM13v2.0 and the maternal haplotype assembly of HG02602.**

The top (chr19_mat-0000001) and bottom (chr19_mat-00000005) tracks show contigs assigned to chr19 in the HG02602 maternal assembly separately. The middle track shows the CHM13v2.0 chr19 reference sequence.

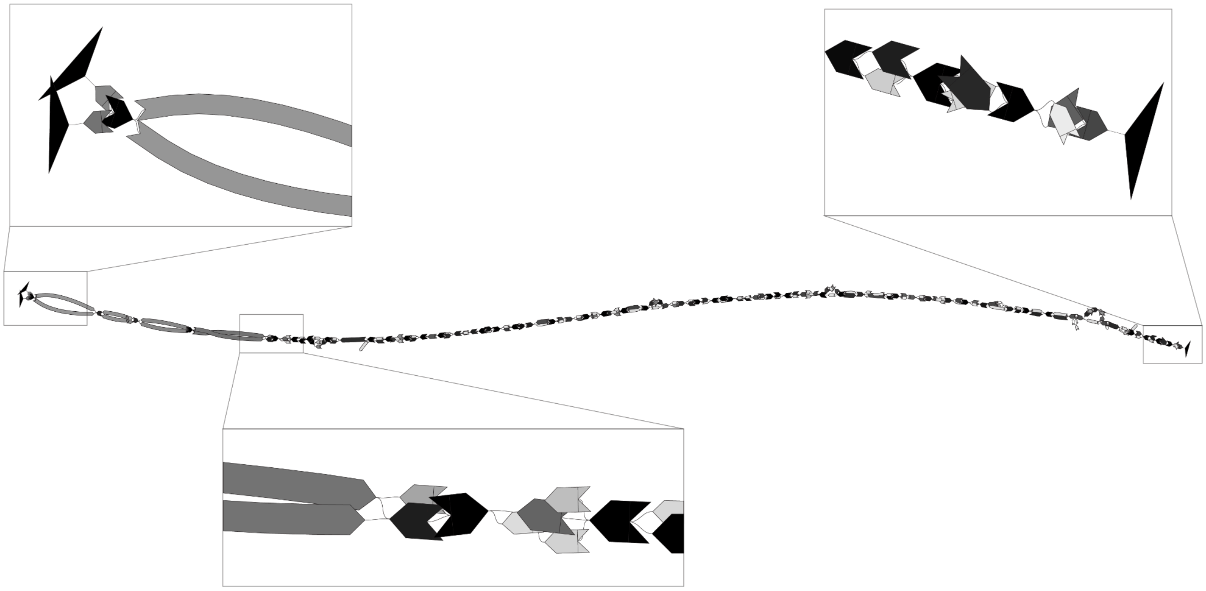

**Supplementary Fig. 18: Diploid assembly graph for chr19 of HG02602**

The diploid assembly graph was visualized using Bandage. The first zoomed-in panel (top left) shows two telomeres at the start of the contig and diploid graph features, with approximately 20× coverage along each branch of the bubble (light gray) and homozygous nodes shown in black. The second zoomed-in panel (bottom) shows the transition from a diploid to a haploid graph structure, accompanied by a twofold increase in coverage (black, approximately 40×) and shorter nodes. The final zoomed-in panel (top right) shows the end of the contig, with a single telomere (black triangle) and diploid-level coverage consistent with the collapse of both haplotypes into a single graph path.

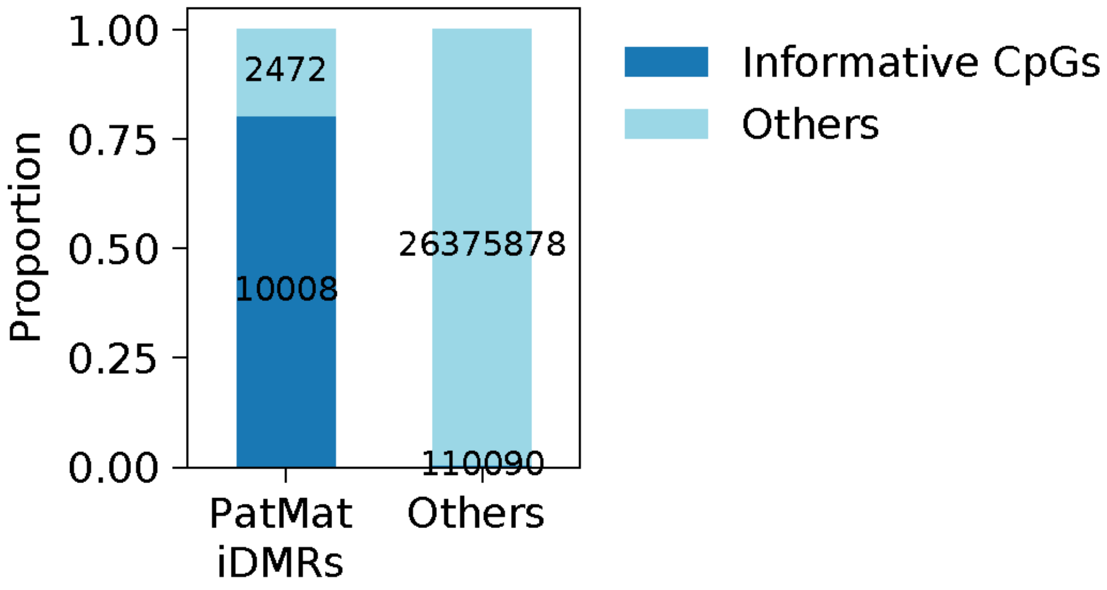


**Supplementary Fig. 19: Enrichment of informative CpGs within PatMat iDMRs.**

Stacked bar plots show the proportions of informative and other CpGs within and outside PatMat iDMRs. Numbers indicate the CpG counts in each category, highlighting the enrichment of informative CpGs within PatMat iDMRs.

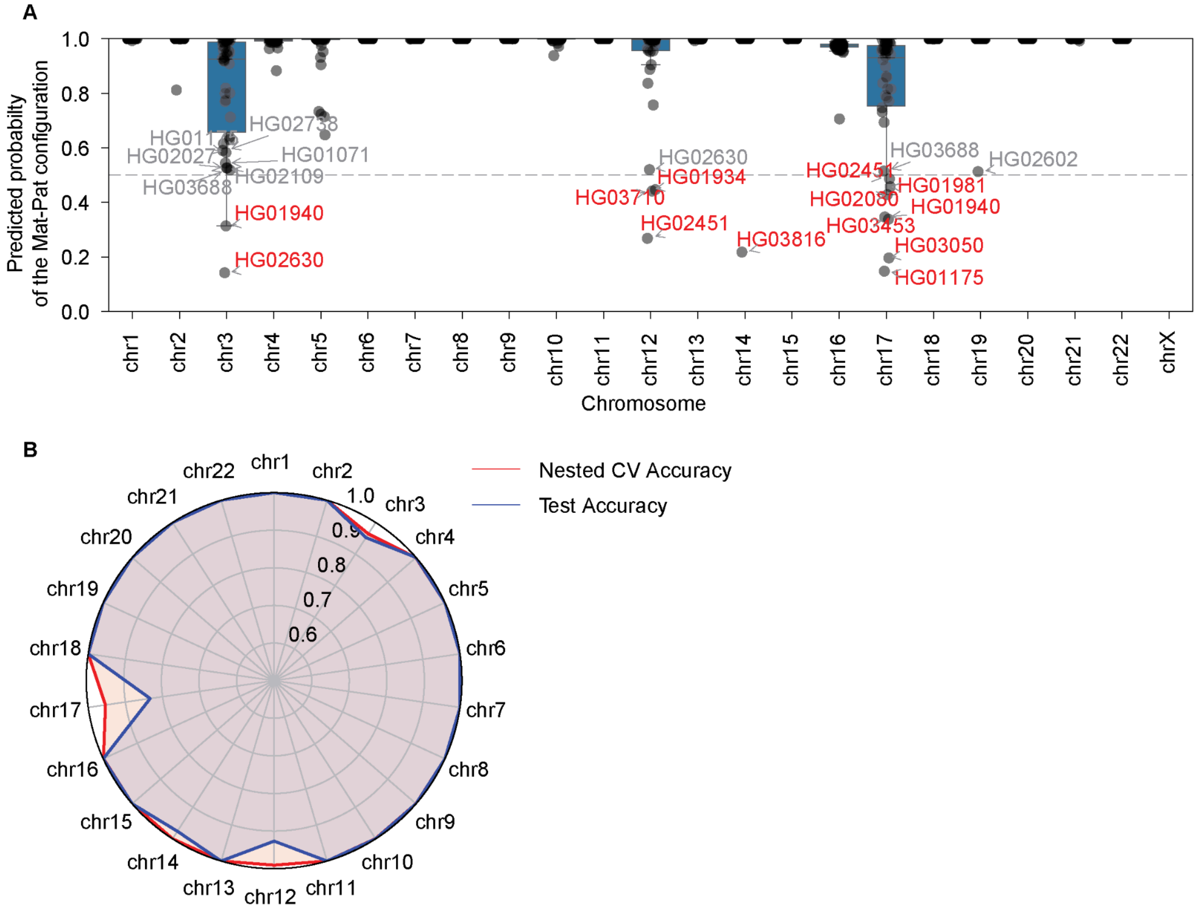


**Supplementary Fig. 20: Model performance using CpGs within PatMat iDMRs**

(A) Predicted probability of the Mat–Pat direction for test samples using a model trained on CpGs within PatMat iDMRs. (B) Model accuracy from nested cross-validation and on the test set.

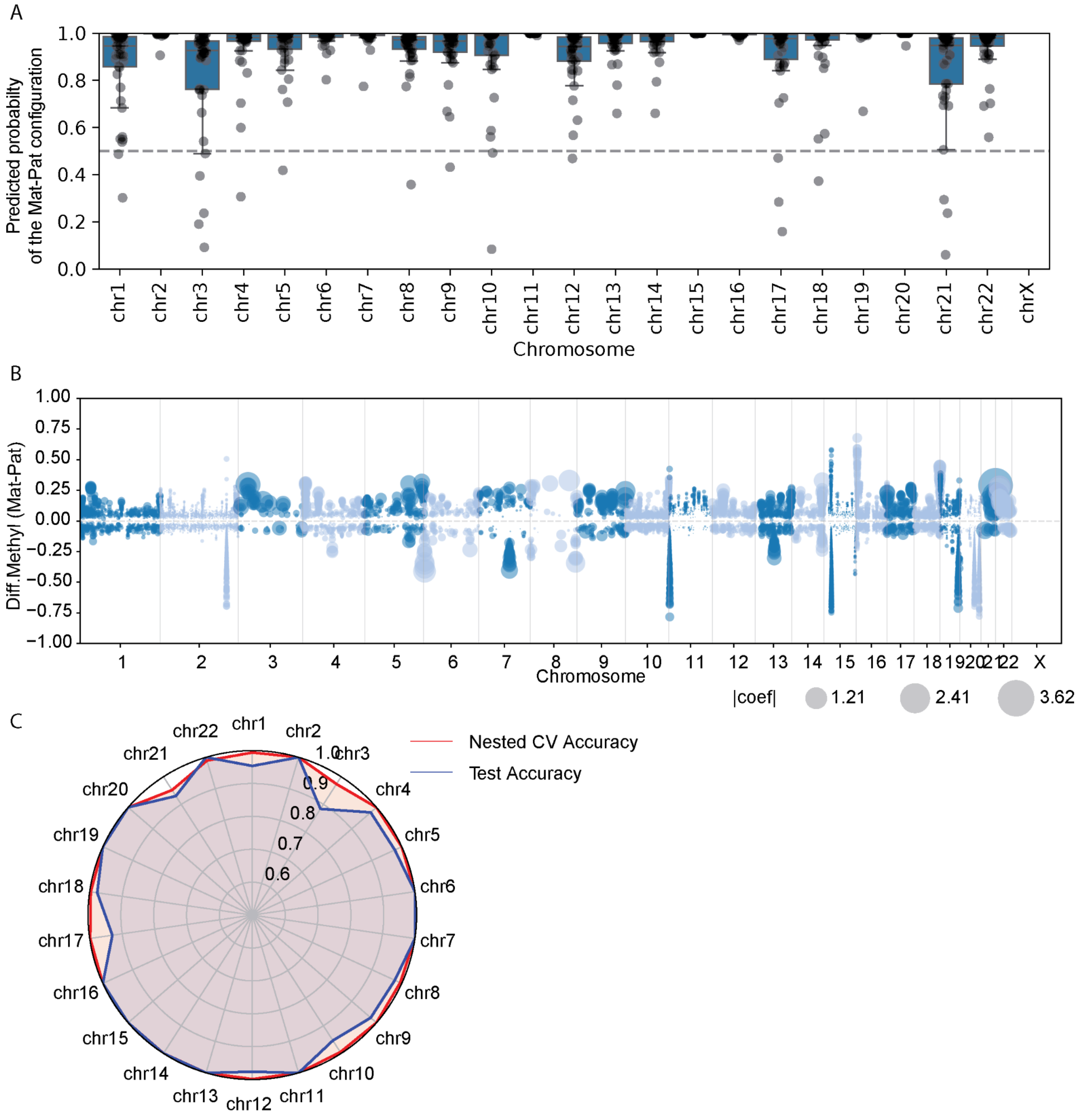


**Supplementary Fig. 21: Performance of a model trained using CpGs outside PatMat iDMRs with a 1-bp padding region**

(A) Predicted probability of the Mat–Pat direction for test samples using a model trained on CpGs outside PatMat iDMRs, with a 1-bp padding region around each iDMR excluded. (B) Differential methylation (Diff.Methy; Mat − Pat) across chromosomes. Point size indicates the absolute value of the learned model coefficient. (C) Model accuracy from nested cross-validation and on the test set.

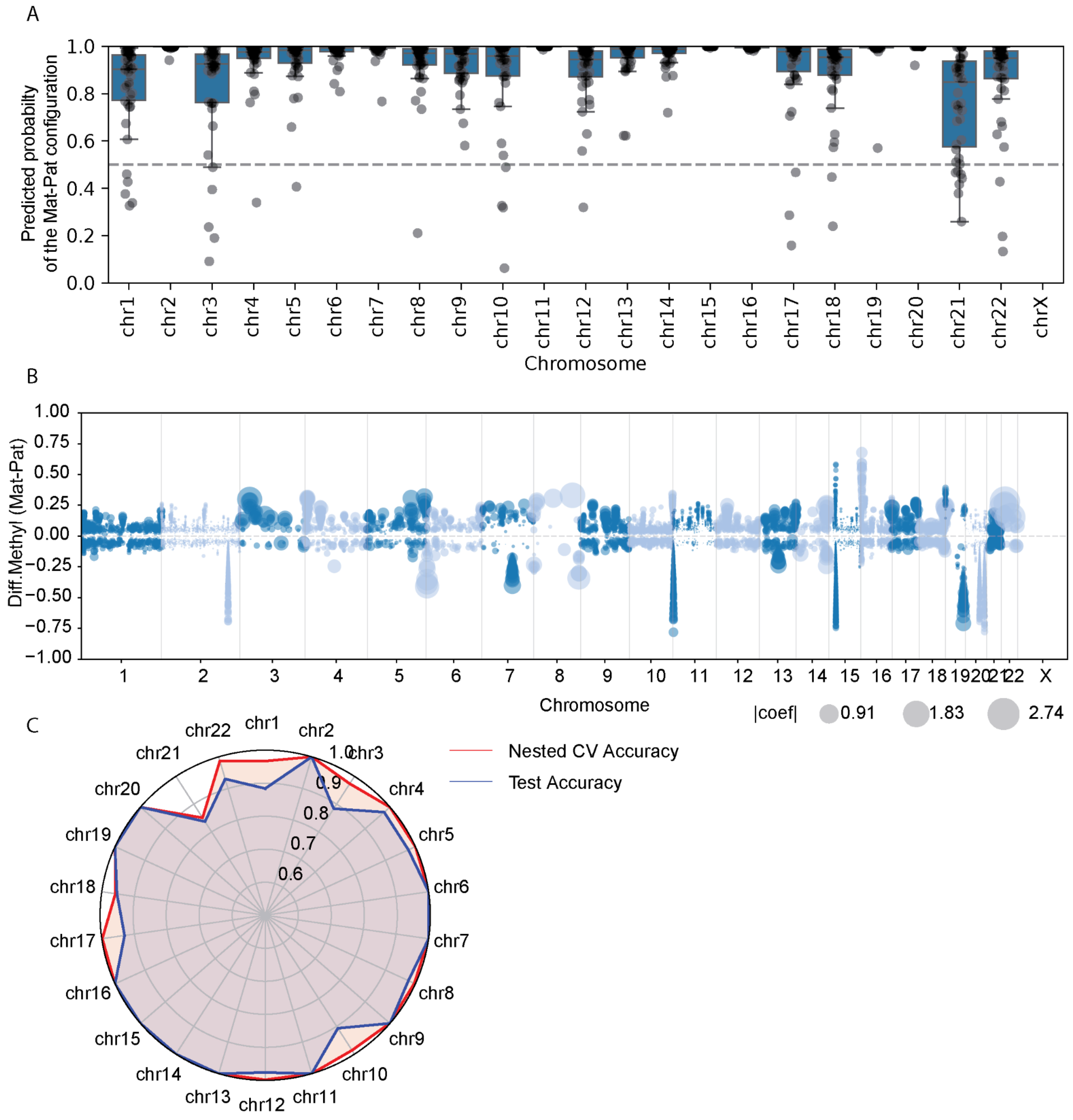


**Supplementary Fig. 22: Performance of a model trained using CpGs outside PatMat iDMRs with a 2-kb padding region**

(A) Predicted probability of the Mat–Pat direction for test samples using a model trained on CpGs outside PatMat iDMRs, with a 2-kb padding region around each iDMR excluded. (B) Differential methylation (Diff.Methy; Mat − Pat) across chromosomes. Point size indicates the absolute value of the learned model coefficient. (C) Model accuracy from nested cross-validation and on the test set.

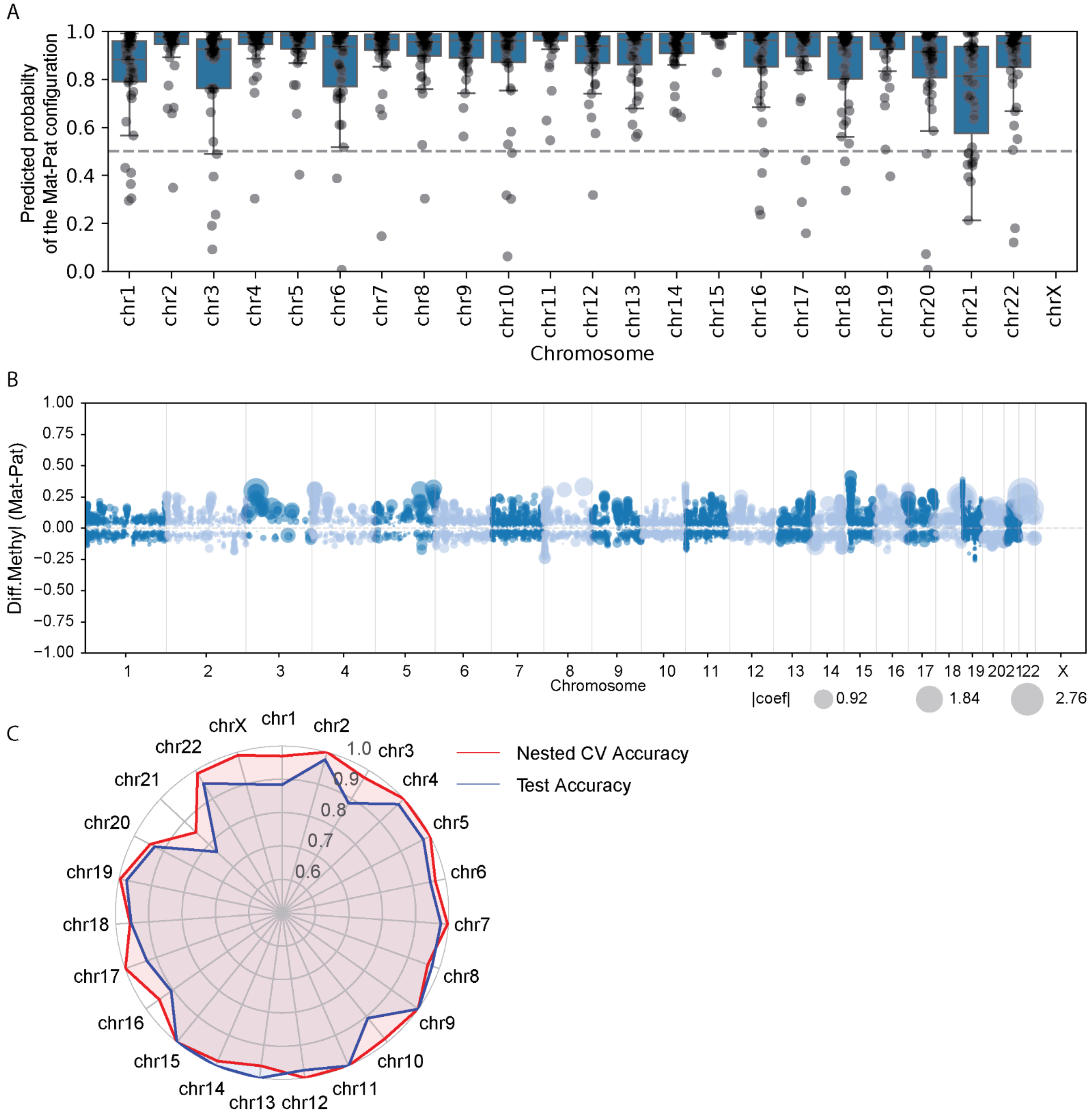


**Supplementary Fig. 23: Performance of a model trained using CpGs outside PatMat iDMRs with a 100-kb padding region**

(A) Predicted probability of the Mat–Pat direction for test samples using a model trained on CpGs outside PatMat iDMRs, with a 100-kb padding region around each iDMR excluded. (B) Differential methylation (Diff.Methy; Mat − Pat) across chromosomes. Point size indicates the absolute value of the learned model coefficient. (C) Model accuracy from nested cross-validation and on the test set.

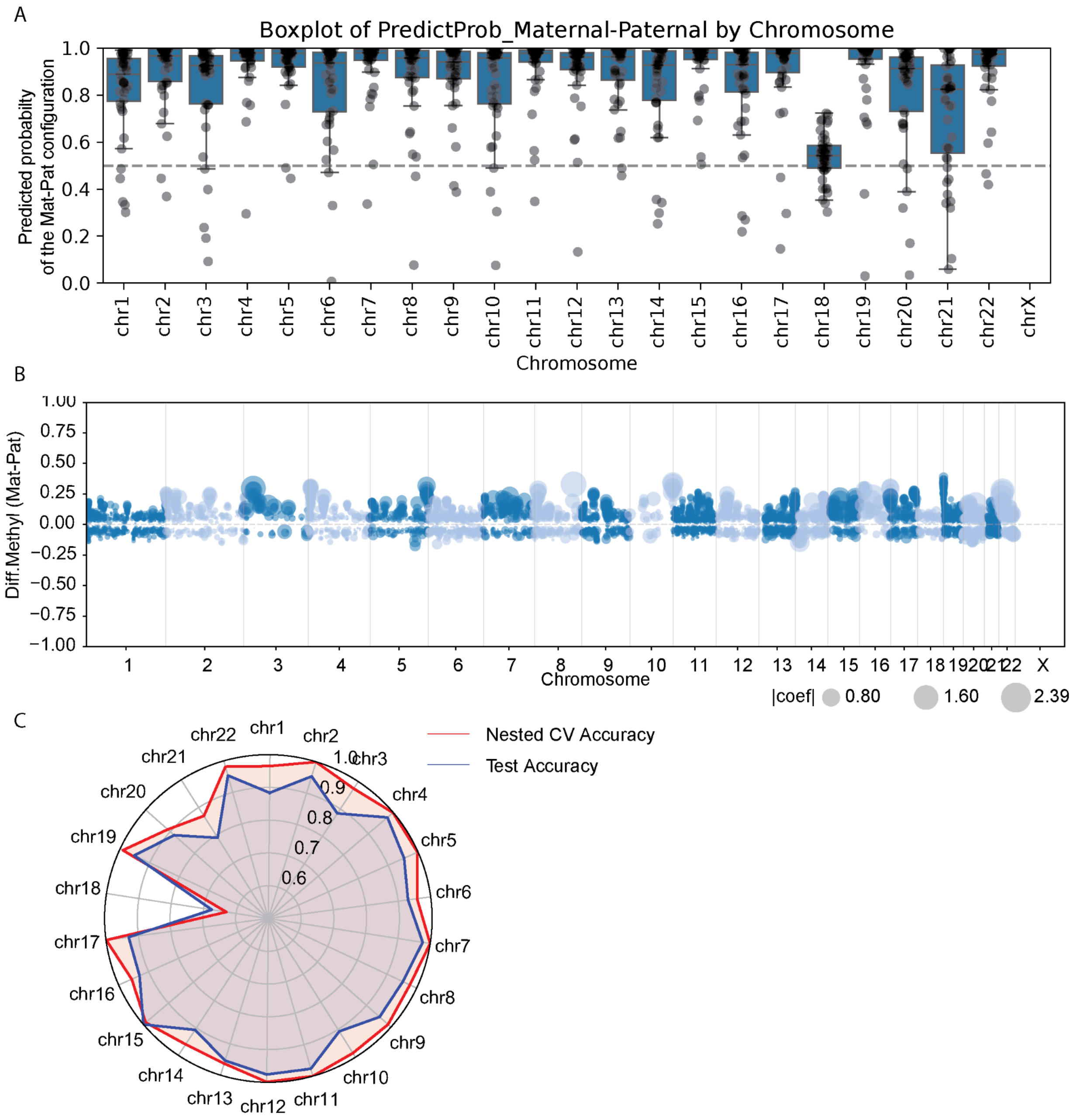


**Supplementary Fig. 24: Performance of a model trained using CpGs outside PatMat iDMRs with a 1 Mb padding region**

(A) Predicted probability of the Mat–Pat direction for test samples using a model trained on CpGs outside PatMat iDMRs, with a 1 Mb padding region around each iDMR excluded. (B) Differential methylation (Diff.Methy; Mat − Pat) across chromosomes. Point size indicates the absolute value of the learned model coefficient. (C) Model accuracy from nested cross-validation and on the test set.

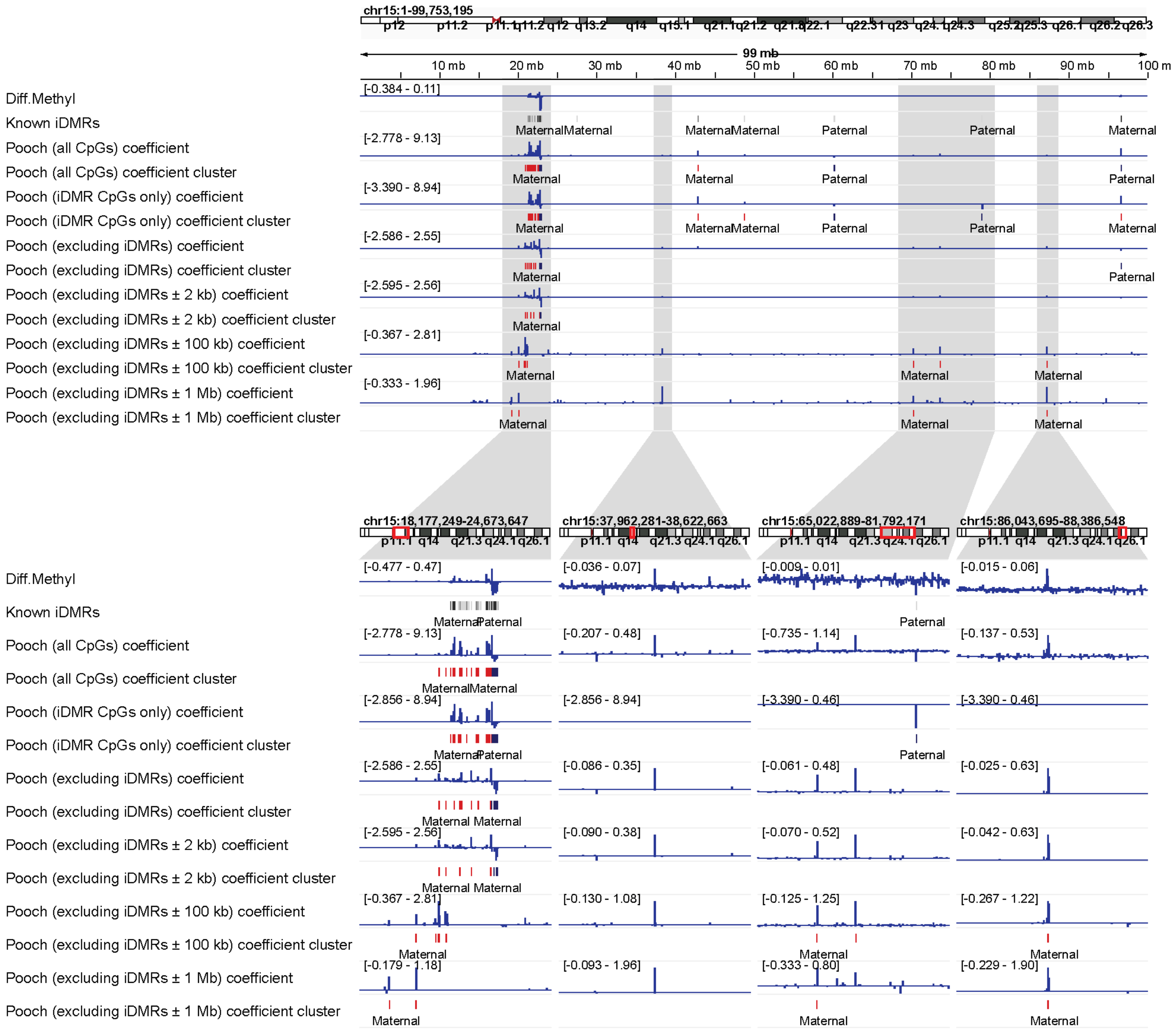


**Supplementary Fig. 25: Overview and close-up genomic plots of chromosome 15 showing coefficients from models trained using different CpG sets defined by the inclusion or exclusion of PatMat iDMRs and their flanking regions.**

Top: chromosome-wide overview. Bottom: close-up views of the highlighted regions. Diff. Methyl indicates maternal minus paternal methylation levels, and PatMat iDMRs indicates previously reported iDMRs. Coefficients from six models and clusters of CpGs with nonzero coefficients are shown. Elastic net coefficients were normalized separately for each model by dividing each coefficient by the maximum absolute coefficient, scaling the largest absolute coefficient to one. The normalized coefficients were then summed within 100-bp genomic bins. Pooch (all CpGs) was trained using all CpGs, whereas Pooch (iDMR CpGs only) was trained using only CpGs within PatMat iDMRs. The remaining four models excluded CpGs within PatMat iDMRs and their flanking regions extending 1 bp, 2 kb, 100 kb, or 1 Mb on either side.

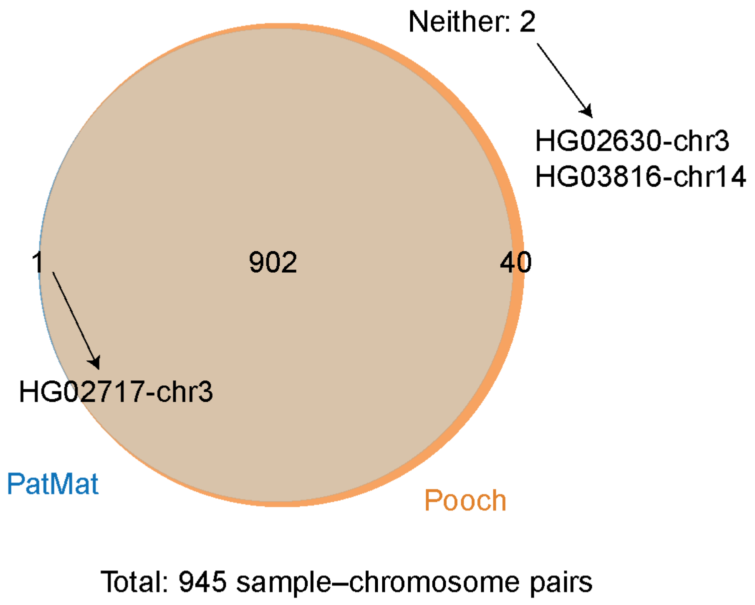


**Supplementary Fig. 26: Comparison of correct parent-of-origin assignments by PatMat and Pooch.**

Venn diagram showing the overlap of correctly classified sample–chromosome pairs among 945 autosomal pairs in the held-out test set. Both methods correctly classified 902 pairs, whereas 40 were correctly classified only by Pooch and one only by PatMat. Two pairs were correctly classified by neither method.

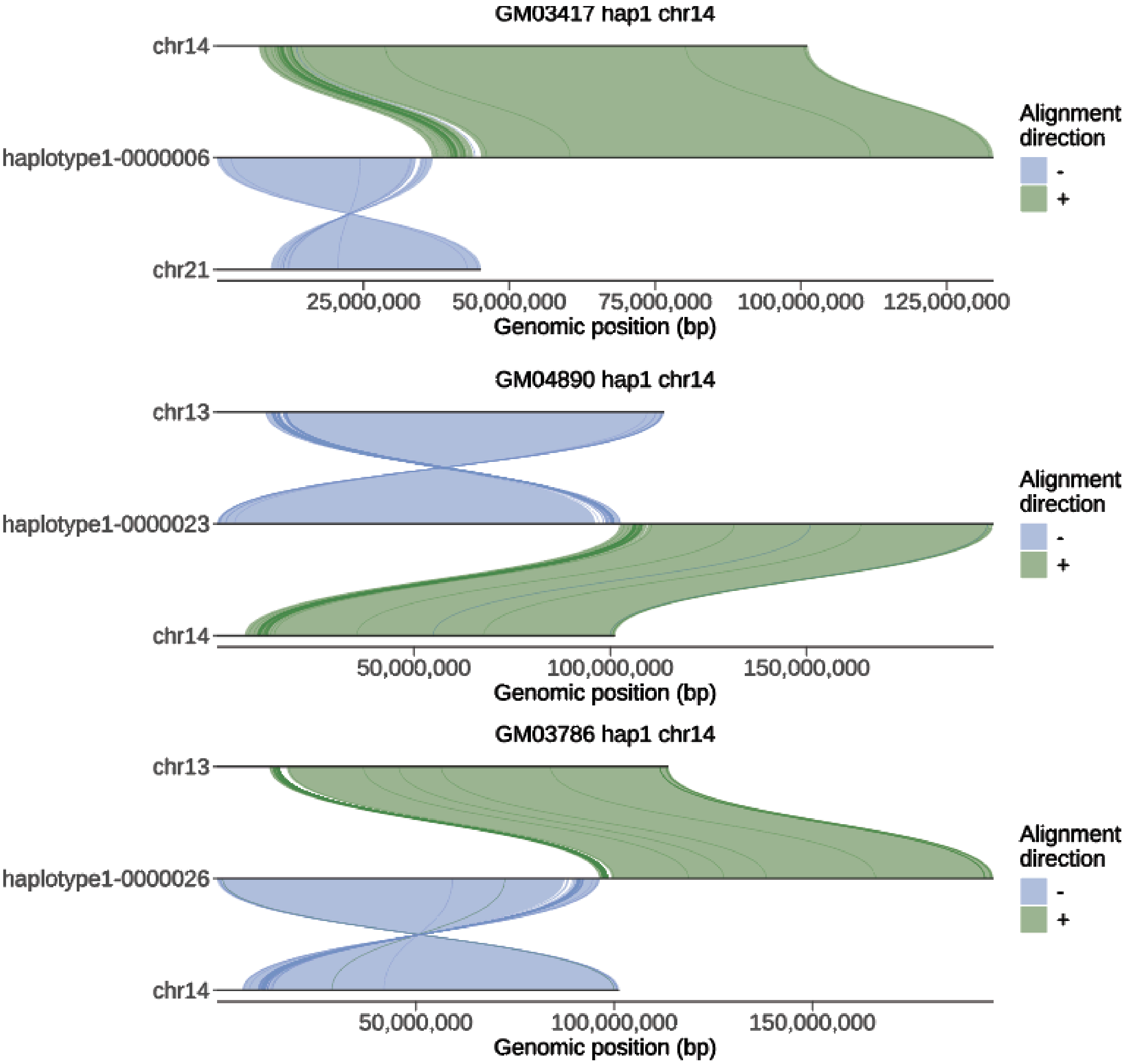


**Supplementary Fig. 27: Syntenic plot of ROB chromosomes**

Synteny plots of the ROB fusion chromosome against the corresponding two chromosomes in the CHM13 reference genome for four ROB carrier samples.

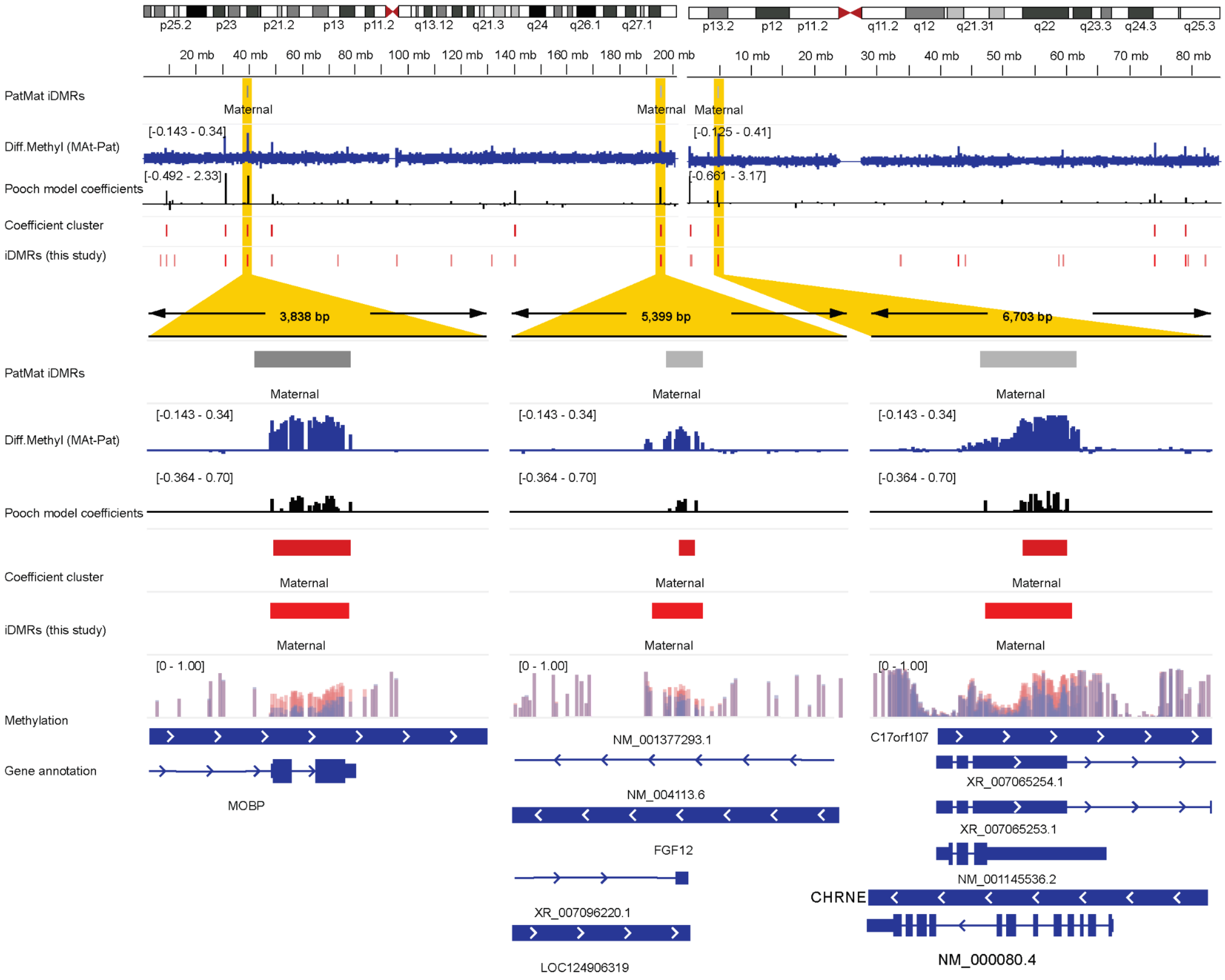


**Supplementary Fig. 28: Overlap of model coefficient clusters and iDMRs identified in this study with PatMat iDMRs on chr3 and chr17.**

The top panels show the same genomic views as Fig. 4A, with PatMat iDMR regions highlighted.

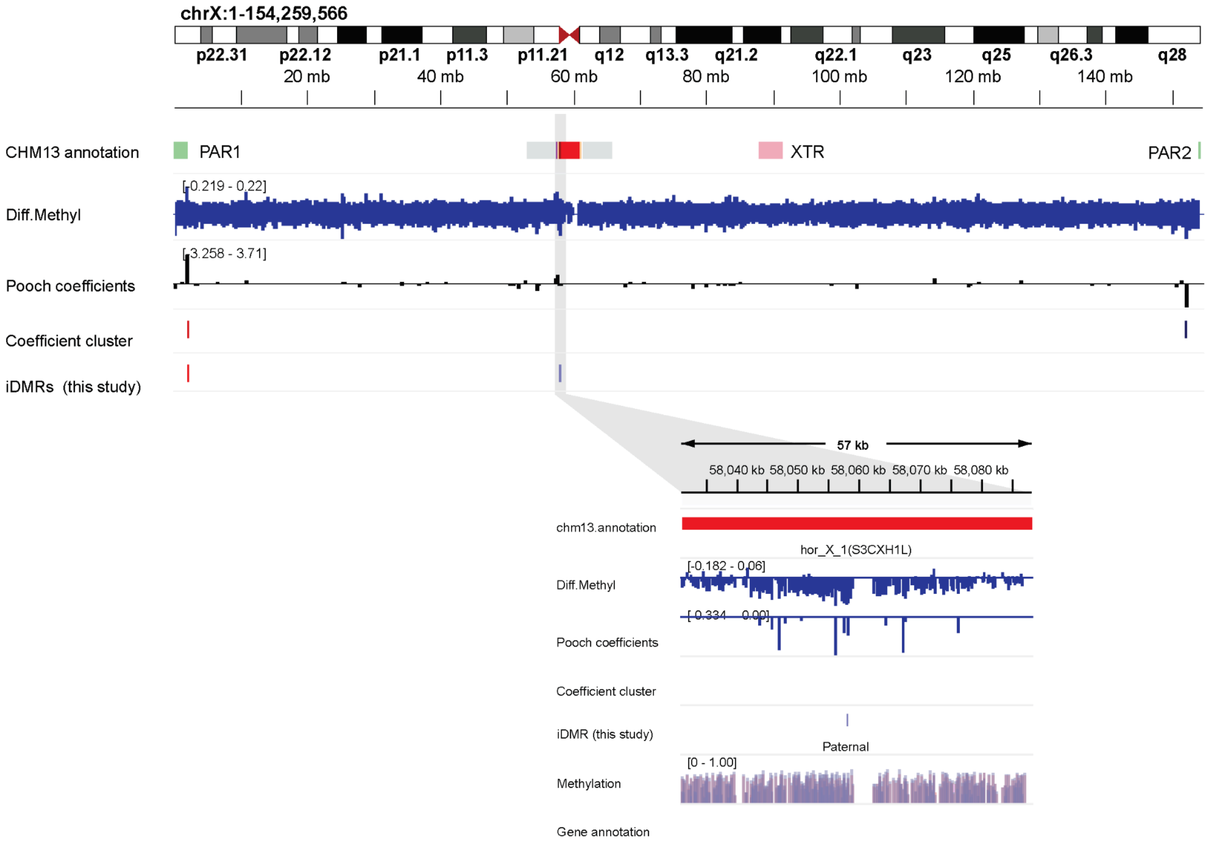


**Supplementary Fig. 29: Zoomed-in genomic views of regions containing iDMRs identified in this study on chrX.**

The top panel is identical to Fig. 4B. The remaining panels show enlarged views of all regions except the one highlighted in Fig. 4C.

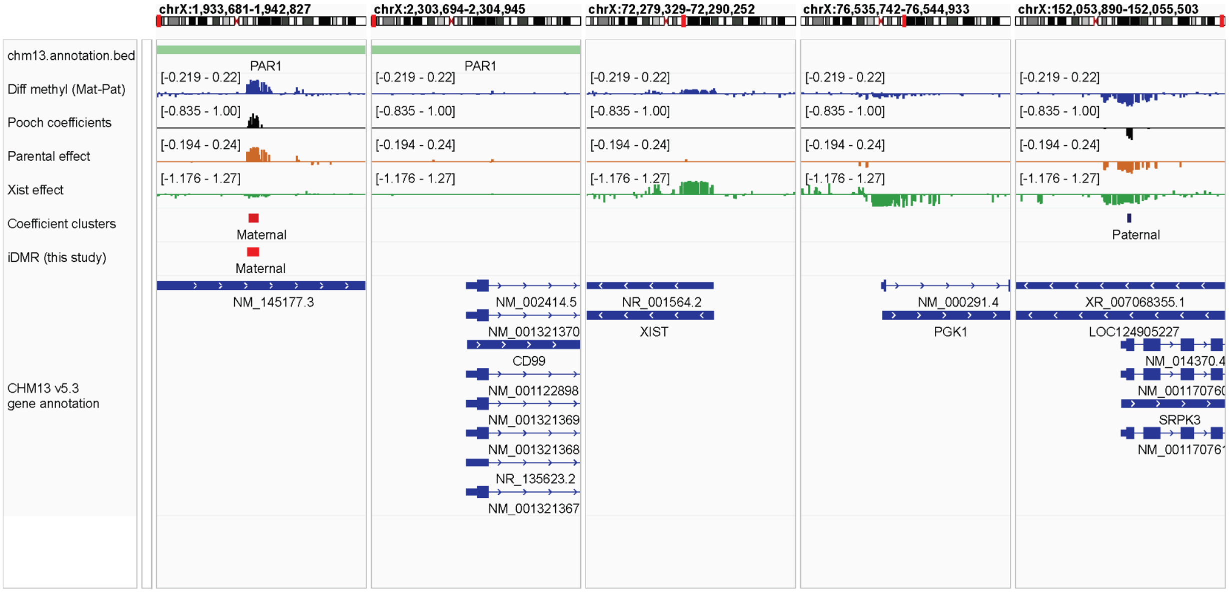


**Supplementary Fig. 30: Zoomed-in genomic views of selected iDMRs and gene promoters on chromosome X.**

The plots show the strong iDMR within an intron of DHRSX in PAR1, *CD99* (an XCI escape gene), *XIST* (a key regulator of XCI), *PGK1* (a known XCI target), and *SRPK3* (a candidate imprinted gene). From top to bottom, the tracks display T2T-CHM13 chromosome annotations, including PAR1, mean differential methylation between parental haplotypes (Mat − Pat), Pooch model coefficients, estimated parental haplotype and *XIST* methylation effects at individual CpG sites during model training, clusters of CpGs with large Pooch model coefficients, iDMRs identified in this study, and gene annotations.

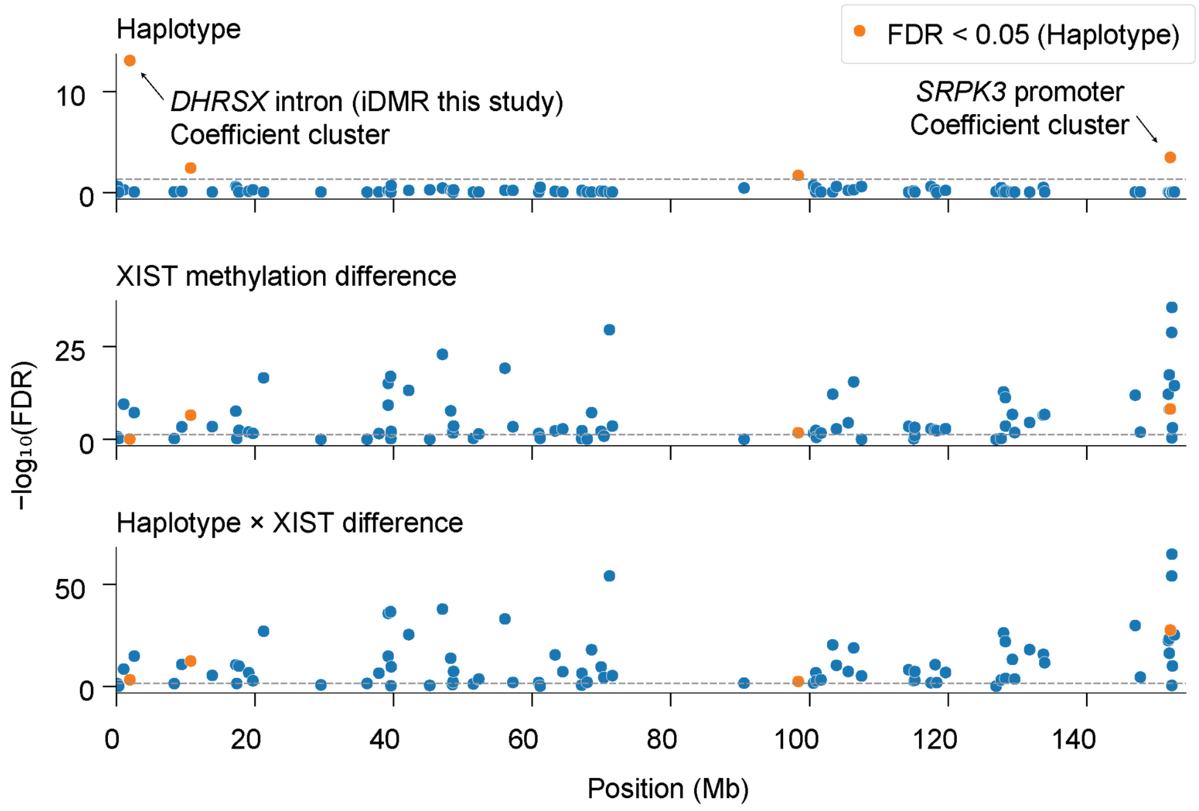


**Supplementary Fig. 31: Interaction analysis of parental haplotype and XIST promoter methylation at previously reported putative imprinting control regions on chromosome X.**

Manhattan plots of interaction analysis results for parental haplotype (top), *XIST* promoter differential methylation (middle), and their interaction (bottom), using previously reported putative ICRs on chromosome X.

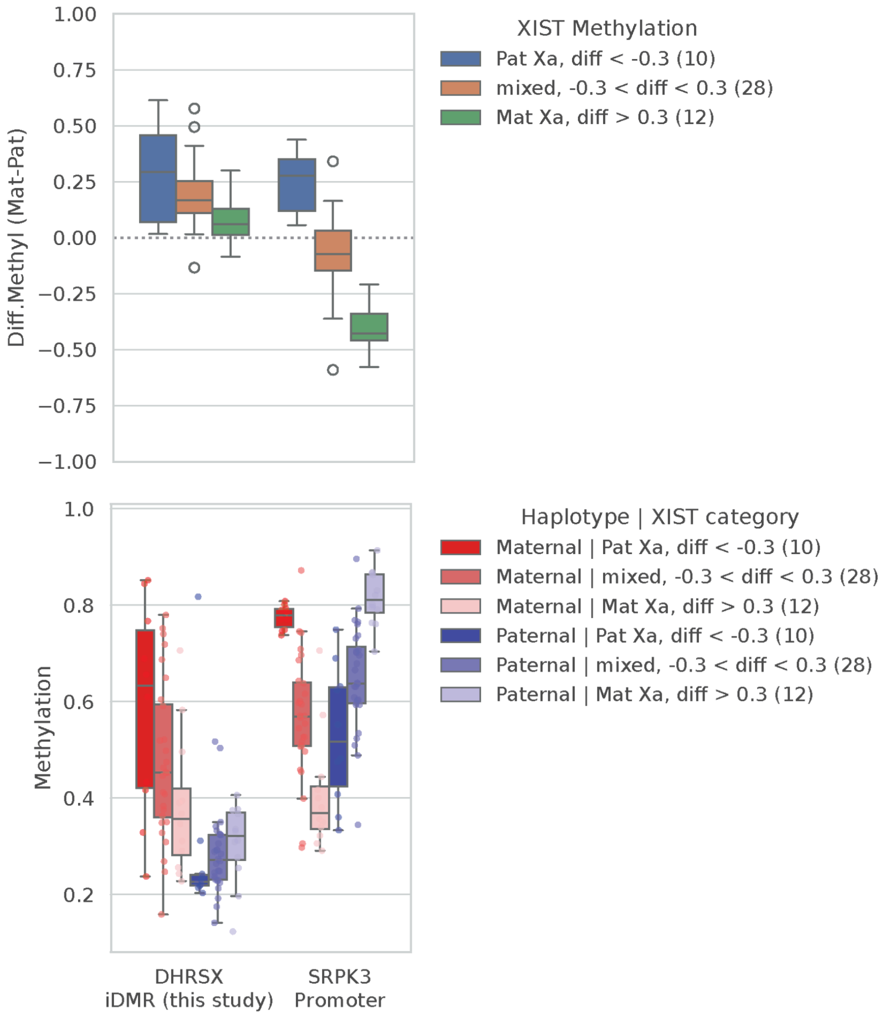


**Supplementary Fig. 32: Differential and mean methylation levels at selected regions and the SRPK3 promoter stratified by XCI status.**

Differential and mean methylation levels are shown for the iDMR in the intron of *DHRSX* and the SRPK3 promoter. Female samples were classified into three groups based on XIST promoter differential methylation (Mat − Pat), with maternal X active (Mat Xa) for values greater than 0.3, paternal X active (Pat Xa) for values below −0.3, and mixed XCI for values between −0.3 and 0.3.

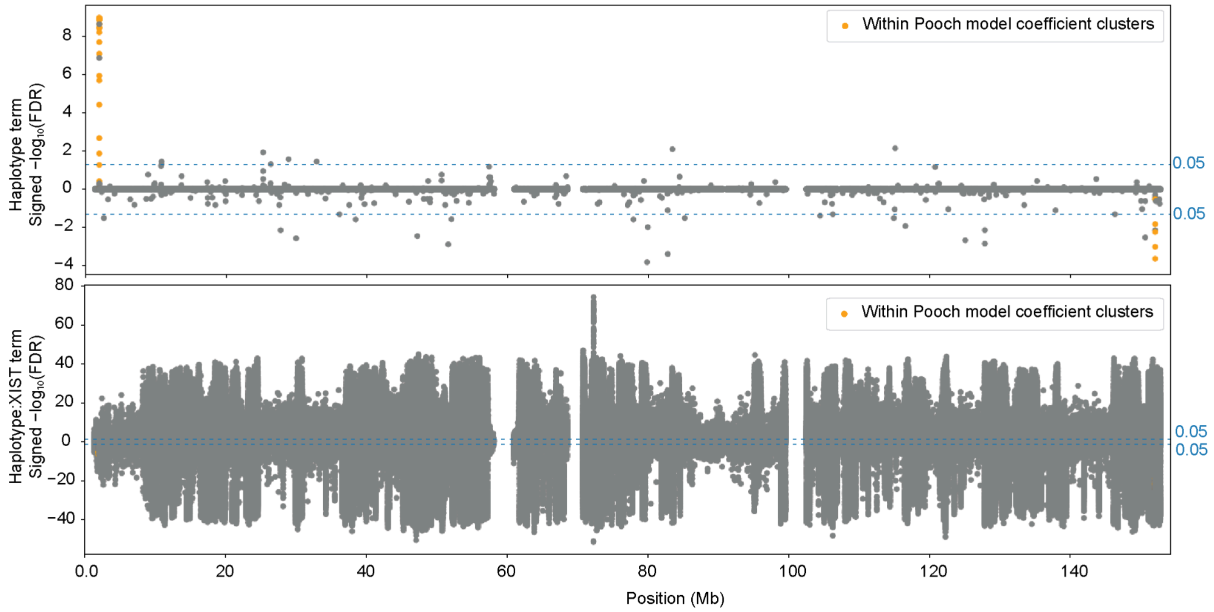


**Supplementary Fig. 33: Manhattan plots of chromosome X-wide CpG-level interaction analysis incorporating parental haplotype and XIST promoter differential methylation.**

The top and bottom panels show signed $-\log_{10}$(FDR) values for the haplotype main effect and haplotype-by-XIST interaction effect, respectively. CpGs within Pooch model coefficient clusters are highlighted in orange, while all other CpGs are shown in gray. Blue horizontal lines indicate the significance threshold of FDR = 0.05 in both positive and negative directions.

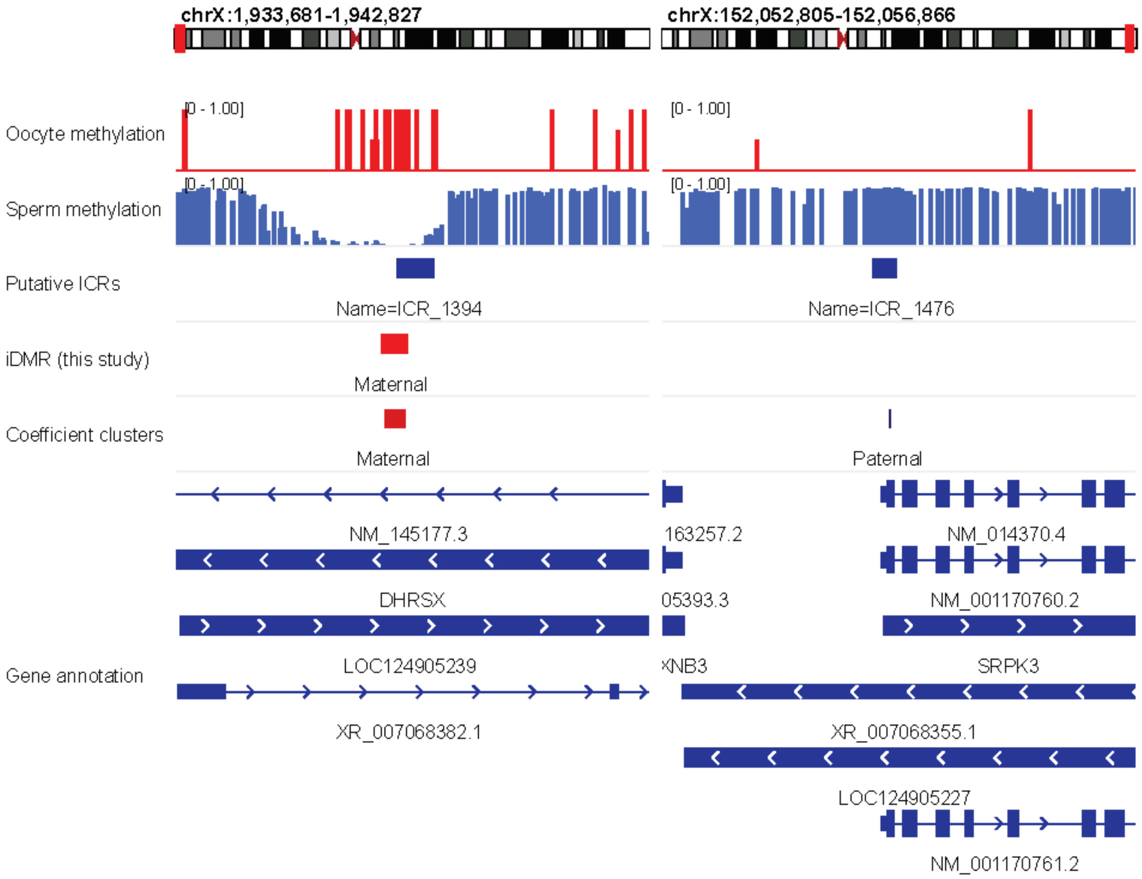


**Supplementary Fig. 34: Oocyte and sperm DNA methylation patterns at the DHRSX and SRPK3 loci.**

Oocyte (Oocyte_100) and sperm (sperm_with_sue_100) methylation data, along with previously reported putative ICRs, were obtained from humanicr.org and lifted over to T2T-CHM13 coordinates. The tracks also display iDMRs identified in this study, clusters of large model coefficients from the Pooch chromosome X model, and gene annotations.

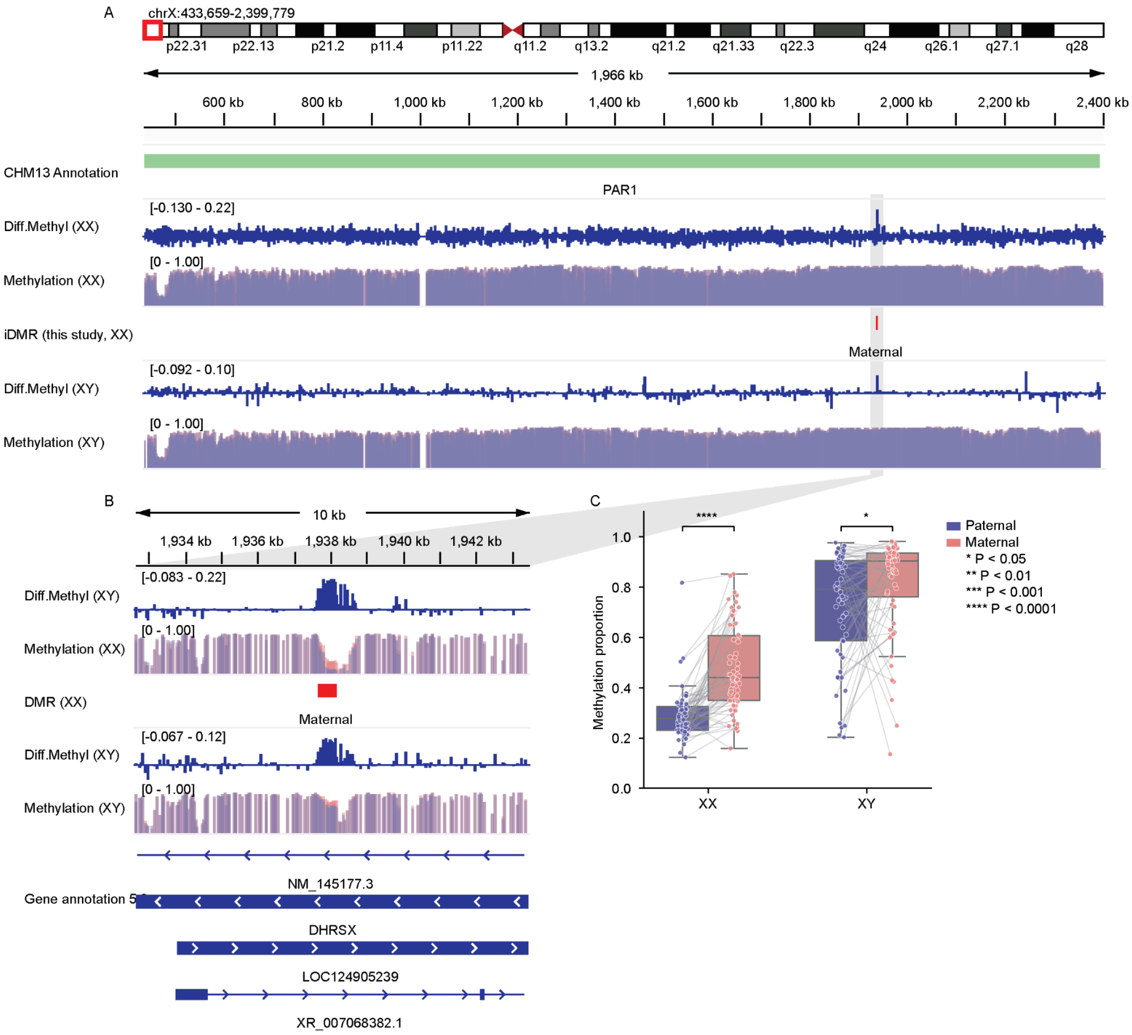


**Supplementary Fig. 35: Genomic comparison of methylation patterns in PAR1 between XX and XY samples.**

(A) Genomic overview of PAR1 with T2T-CHM13 annotations. (B) Zoomed-in view of the *DHRSX* region showing differential methylation, mean methylation levels, and iDMRs in XX samples, following the same format used throughout the study. The "Diff. methylation (XY)" track displays methylation differences between the maternally inherited X chromosome (mX) and the paternally inherited Y chromosome lifted over to chromosome X coordinates (pseudo-pX), calculated as mX − pseudo-pX. The "Methylation (XY)" track overlays the mean methylation levels of mX (pink) and pseudo-pX (blue). (C) Comparison of mean methylation levels between maternal and paternal haplotypes in XX (mX and pX) and XY (mX and pseudo-pX) samples. P-values were calculated using paired t-tests, with significance levels indicated above the box plots. All 59 XX samples and 63 of 64 XY samples were included, with HG02514 excluded due to the absence of maternal X chromosome CpG methylation calls within the analyzed region.

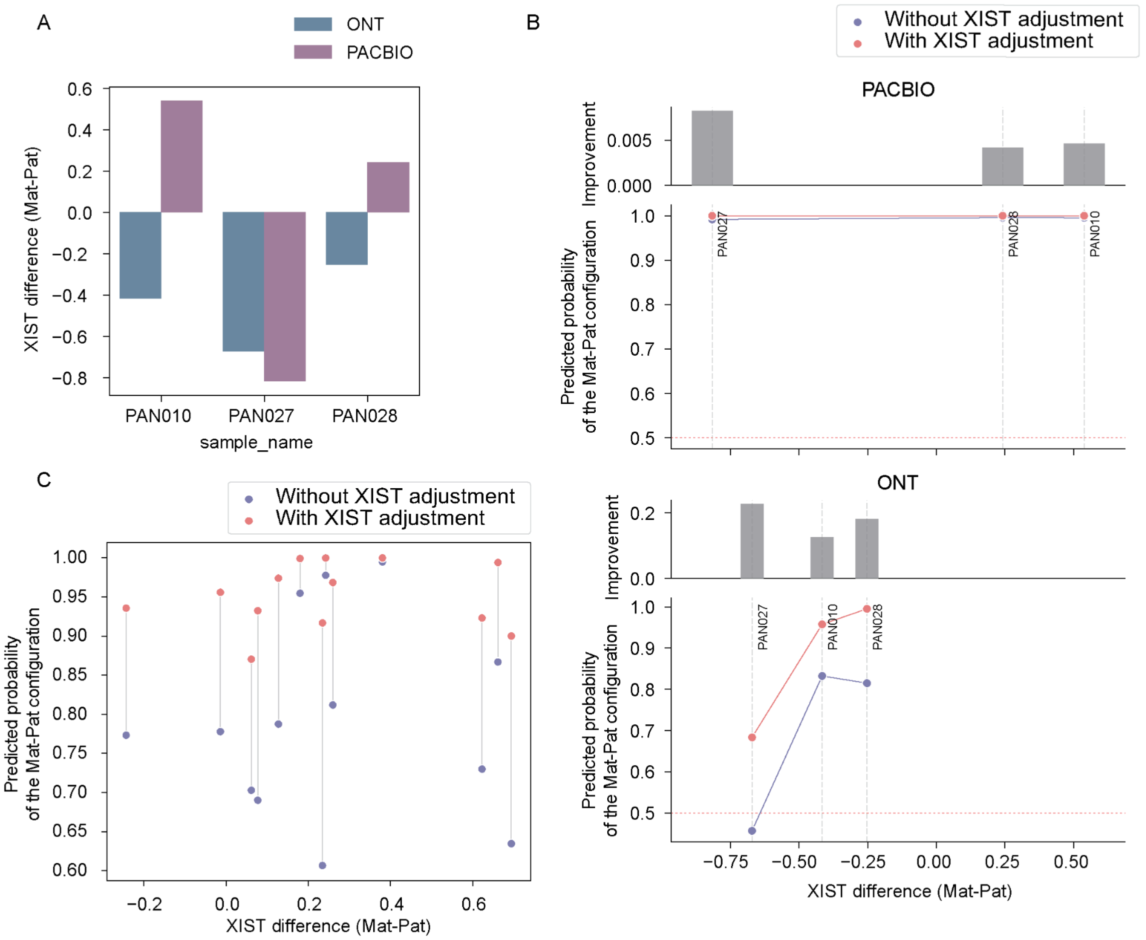


**Supplementary Fig. 36: Comparison of Pooch model performance on chromosome X with and without adjustment for XIST promoter differential methylation.**

(A) XIST promoter differential methylation in the T2T pedigree, shown by individual and sequencing platform. Predicted probabilities of the Mat − Pat configuration from models with and without XIST adjustment are shown for (B) T2T pedigree samples, stratified by sequencing platform (top, PacBio; bottom, ONT), and (C) HPRC test samples. Lines in (C) connect predictions from the two models for the same individual.
